# Multiple broadly neutralizing antibody lineages can co-exist and mature in the same germinal centres

**DOI:** 10.1101/2025.11.09.687328

**Authors:** Amar Kumar Garg, Sebastian C. Binder, Michael Meyer-Hermann

## Abstract

An efficacious HIV vaccine will need to generate broadly neutralizing antibodies (bnAbs) against distinct viral epitopes. To facilitate this, immunogens targeting precursor B cells of bnAbs have been developed. With this strategy, individual immunogens can even target multiple lineages, thereby beneficially limiting the number of immunogens needed for a multi-bnAb generating vaccine. However, it is unclear whether this diminishes responses compared to isolated targeting of lineages with distinct immunogens. Here, we address this using an *in silico* model of naive B cell activation and affinity maturation in germinal centres. By incorporating (i) precursor properties and (ii) epitope masking by antibodies obtained from germinal centre-derived plasma cells, the model recapitulated features of bnAb lineage evolution as seen in pre-clinical mouse models. Subsequent model analysis suggested that under physiologically relevant conditions, priming of multiple bnAb lineages with a single immunogen was additive, thus, supporting the development of vaccines that target multiple lineages.

## INTRODUCTION

Despite significant efforts spanning over 3 decades, we still lack an effective vaccine against HIV, majorly because of the high variability of HIV’s envelope protein (ENV) that arises due to the virus’ mutability^1,2^. In this context, vaccine-mediated generation of broadly neutralizing antibodies (bnAbs) is an attractive prospect as these antibodies can target diverse HIV strains by binding to conserved sites of the ENV protein^3–5^. However, in natural infections, bnAbs emerge in a small percentage of individuals, over several months to years and require extensive somatic hypermutation (SHM)^3,5–8^. Unsurprisingly, producing bnAbs via vaccination has proven to be an immense challenge and has not been possible so far.

A major bottleneck for vaccine-mediated elicitation of bnAbs is the activation and expansion of their precursor B cells. Naive B cells with the potential to evolve and generate bnAbs are rare in humans, i.e., a range of few bnAb precursor B cells per million naive B cells or even lower^9–11^. Moreover, these cells typically exhibit low or undetectable binding to ENV proteins derived from commonly circulating viruses^12–15^, making their activation difficult.

To address this issue, one strategy has been to employ germline-targeting immunogens. These are ENV-based antigens engineered to bind bnAb precursor B cells, also known as germline precursors, with high affinity to facilitate their activation and recruitment to the germinal centre (GC). Within GCs, these precursor B cells can then acquire a limited number of early mutations that are relevant for the bnAb lineage and enable the production of corresponding memory cells^11,16–20^. This combination of activation and maturation of precursor B cells in GCs is subsequently referred to as “priming”. Germline-targeting immunogens have been successfully used for priming bnAb lineages in animal models^11,16–18,21–23^ and in humans^19^. Notably, post priming, boosting with other suitable immunogens is still required to accumulate additional mutations needed to achieve broad neutralization^3,7,8,23–29^.

Strategies targeting single bnAb lineages may not be enough, as demonstrated by the recent Antibody Mediated Protection trials. In these studies, passive immunization with the bnAb VRC01 did not significantly improve protection against HIV transmission^30^. However, the study noted reduced transmission of HIV viruses sensitive to VRC01, which suggests that a protective vaccine requires multiple bnAbs^31–33^. This can be challenging as with current approaches, numerous immunogens are already needed to elicit individual bnAbs^3,7,8,32^. At least at the priming stage, this obstacle can be overcome by employing immunogens that can simultaneously prime multiple bnAb lineages. Although germline-targeting studies are often performed with emphasis on closely related precursor B cells^13,16–18,21,22^, advanced immunogens such as eOD-GT8 and GT1, and their variants have improved affinities for precursor B cells of distinct bnAbs^10,20,34^. It is not known whether simultaneously targeting multiple bnAb lineages with a single antigen can diminish individual lineage output when compared to targeting individual lineages with distinct antigens.

Germline-targeting also stimulates B cells that lack features commonly attributed to bnAb precursor B cells, such as the restricted usage of specific V gene segments like VH1-2 for VRC01 bnAb precursors or the often seen long heavy chain complementarity determining region 3^3,10,16,19,23,35–37^. Thus, concurrent priming of diverse lineages will invoke competition with cells unlikely to develop into bnAb-producing cells. A comprehensive understanding of this interlineage competition is lacking and is an important knowledge gap. Illuminating this will be useful in assessing the utility of immunogens that target multiple bnAb lineages in parallel.

To address the competition between bnAb and non-bnAb precursor B cells and their respective progeny in GC responses (“bnAb-” and “non-bnAb-type GC B cells”), we mechanistically considered the evolution of B cells (Fig.1). Post immunization, naive B cells in secondary lymph nodes are activated via acquisition of antigen and other signals such as those delivered by CD4 T cells^38–40^. These activated cells can then migrate to the B cell follicle and proliferate to establish GCs^41,42^. Within GCs, B cells compete against each other for survival, which is dictated by the acquisition of antigen and T follicular helper (Tfh) cell signals. Importantly, cells that acquire higher amounts of antigen are more likely to receive Tfh signals^43–45^. Post this, B cells proliferate with mutations, allowing a minority of cells to acquire higher affinities. The cyclic repetition of selection and proliferation with mutations increases GC B cell affinities over time, also known as affinity maturation^42,46–49^. Thus, B cell evolution in the pre-GC and GC phases is inherently competitive. As a consequence, precursor B cells with higher precursor affinity and/or abundance (frequency) are more likely to prosper in these phases^16–18,22,40,50–53^.

**Figure 1.**
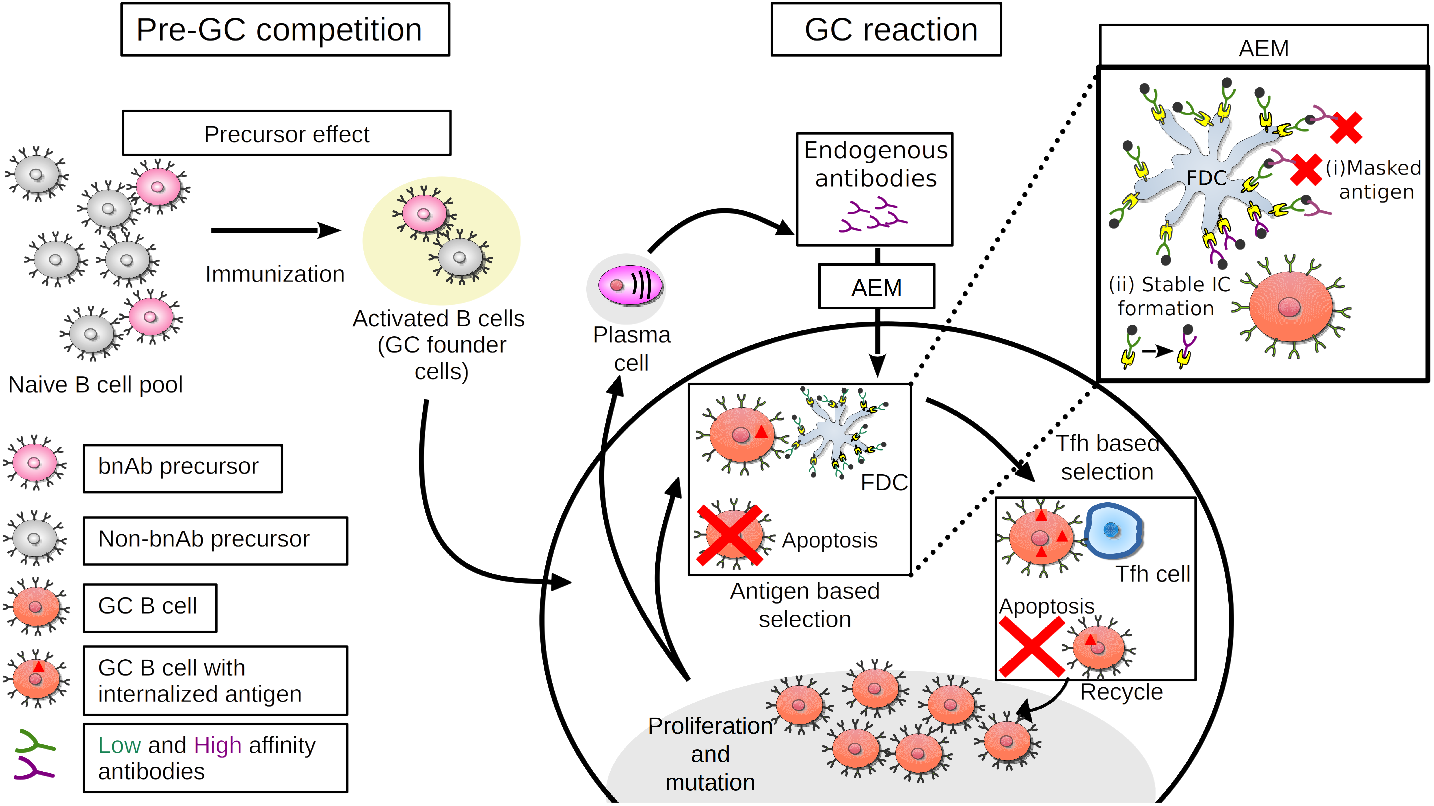
Schematic of *in silico* B cell responses post immunization. Precursor B cells get activated upon immunization and seed GCs as founder cells. The fraction of bnAb- and non-bnAb-type founder cells is calculated via Equations 1 and 3 using their respective precursor frequencies and affinities (precursor effect). Within GCs, B cells that acquire antigen from ICs presented on FDCs and Tfh cell signals survive, proliferate, and mutate, thus, resulting in affinity maturation. As a consequence, affinities of GC-derived plasma cells and associated antibodies also increase with time. These higher affinity antibodies can localize within the GC and mediate AEM. This involves (i) direct masking of antigenic epitopes by antibodies and (ii) formation of more stable ICs that reduce the probability of B cell antigen acquisition. Acronyms used: AEM antibody-epitope-masking; FDC - follicular dendritic cell; GC - germinal centre; IC - immune complex and Tfh cell - T follicular helper cell.

Apart from features of precursor cells (precursor effect), B cell responses can also be modulated by circulating antibodies in the serum. These antibodies, sourced from external injections or GC-derived plasma cells, can localize within GCs and modulate antigen presentation to B cells^54^. Antigen within the GC is presented on follicular dendritic cells (FDCs) bound in immune complexes (ICs). Serum antibodies, in particular high affinity ones, can preferentially form Ics on FDCs by replacing existing lower affinity IC-forming antibodies. They may also mask ICs and impede B cell access to antigen. As B cells will now have to acquire antigen from a more stable IC configuration and/or from fewer exposed ICs, this effect of antibody-epitope-masking (AEM) can increase B cell selection stringency^54–56^. Moreover, in multi-epitope environments, AEM driven by epitope-specific pre-existing antibodies can divert GC B cell responses to alternate epitopes^57–61^.

Building on the above observations, we hypothesized that B cell responses in primary GCs, including those of bnAb lineages, are shaped by a combination of the precursor effect and AEM. For primary GCs, AEM would be mainly mediated by endogenous antibodies that are generated by GC-derived plasma cells. As in the later phase of the GC reaction, these antibodies exhibit higher affinity and concentration, substantial suppression of the antibodies’ cognate epitopes can be expected. Consequently, AEM would direct GC B cell responses to alternate, less targeted epitopes. Importantly, antibodies produced by both bnAb- and non-bnAb-type GC B cells can mediate AEM.

To test this hypothesis, we developed an *in silico* model of B cell activation and affinity maturation. Mathematical models have been used before as effective tools to investigate the mechanisms of B cell responses, including GCs and bnAb development^62–72^. A few works concentrated on the simulation of AEM and its impact on modulating affinity maturation and GC dynamics^54,57,73–76^. In this work, by incorporating the precursor effect and AEM, our simulations qualitatively recapitulated the features of bnAb lineage priming as seen in independent experiments^17,18^, thus supporting the hypothesis. Furthermore, we predicted with the model how variations in features of non-bnAb precursor B cells can drive the precursor effect and AEM and impact priming of individual and multiple bnAb lineages under physiologically relevant conditions. Our simulations demarcate scenarios where simultaneous priming of multiple bnAb lineages can be optimal, thus, informing a rational design of germline-targeting immunogens.

## RESULTS

In this work, B cell responses in the pre-GC and GC phases were simulated by employing a mathematical model (Fig.1, see also STAR METHODS). The pre-GC phase model was used to estimate the composition of activated B cells that seed GCs (“founder cells”) based on naive B cell receptor binding to antigen. As B cell activation is competitive, lineages stemming from naive cells with higher precursor affinities and/or frequencies are likely to outcompete others in getting activation signals and establishing GC reactions^40,50,51,53^. Accordingly, we calculated the fraction of founder cells of a bnAb lineage as a function of the frequencies and affinities of antigen-binding bnAb and non-bnAb precursor B cells (Equation 1 and 3). The equations were formulated such that in the lower limit, this fraction increased with increasing frequency and average affinity of bnAb precursor B cells, while saturating when these values reached their upper bounds. Affinities of precursor B cells were averaged as the objective was to track the entire bnAb lineage rather than individual precursor B cells. Moreover, since affinity is inversely related to the dissociation constant (*K*_D_), average precursor B cell affinity was calculated as the harmonic mean of *K*_D_ values. Consistent with experiments, each GC was assumed to be seeded by ∼ 180 − 200 founder cells^77^.

The GC phase was modelled with a spatio-temporal simulation framework where B cells, FDCs and Tfh cells were defined as individual objects with distinct properties^57,64,78^. The model included representations of the GC architecture and zones^42,79^ along with chemokine gradients and B cell motility^80,81^ consistent with two-photon microscopy experiments^45,82–84^. The model also incorporated features of B cell affinity maturation such as selection, mutation and proliferation^42,46–49,57,62,64,85,86^. Masking of antigenic epitopes caused by endogenous antibodies upon their trafficking back to the GC was represented by chemical kinetics and was, thus, dependent on antibody affinity and concentration^54,57^. As these antibodies may form more stable ICs with antigen on FDCs, the reduction in the probability of a B cell to acquire antigen was scaled down in dependence on the average affinity of circulating antibodies^54^. Overall, this complex simulation framework integrated various GC-related biological phenomena and aligned well with experiments, thus offering a realistic representation of associated B cell dynamics.

To initialize the GC reaction, founder cells were tagged as bnAb- or non-bnAb-type, depending on the pre-GC phase calculations. This tag was inherited upon B cell proliferation, thus allowing tracking of these populations. The extent of priming of bnAb lineages was estimated by tracking the percentage and absolute number of bnAb-type GC B cells through the course of the GC reaction.

### Simulations recapitulate the precursor effect

At first, we used the aforementioned setup to examine the role of bnAb precursor B cell affinity and frequency on priming of bnAb lineages. For model validation, experimental data from the work of Huang et al.^17^ was used. In this study, wildtype C57BL/6 mice received one of three distinct precursor B cells of the VRC01 bnAb that were characterized by their affinities (*K*_D_) as high (0.125*µM*), intermediate (1.3*µM*) or low (18.5*µM*). Furthermore, the frequency of transferred precursor B cells was 10-fold serially diluted over four possible values ranging from 1 bnAb precursor in 10^3^ B cells to 1 in 10^6^. These mice were then immunized with the germline-targeting immunogen eOD-GT8 60mer, following which bnAb-type GC B cells were quantified. Importantly, mice with 1 in 10^6^ precursor frequency approximated the physiological setting, as precursor B cells of the VRC01 bnAb having substantial affinity (*K*_D_ *<* 3*µM*) were approximated to be similarly rare in humans^9,10^.

To replicate these conditions *in silico*, we assumed a mono-epitope antigen capable of binding to both bnAb and non-bnAb precursor B cells. Founder cell fractions were estimated using Equations 1 and 3 for the different settings (for parameter values see section Pre-GC Model) and used to simulate GCs. The percentage of bnAb-type GC B cells in simulations (Fig.2A) closely approximated the experimental values, which varied over five orders of magnitude, thus validating our model. The associated temporal trajectories were shown in Fig.S1B-M.

**Figure 2.**
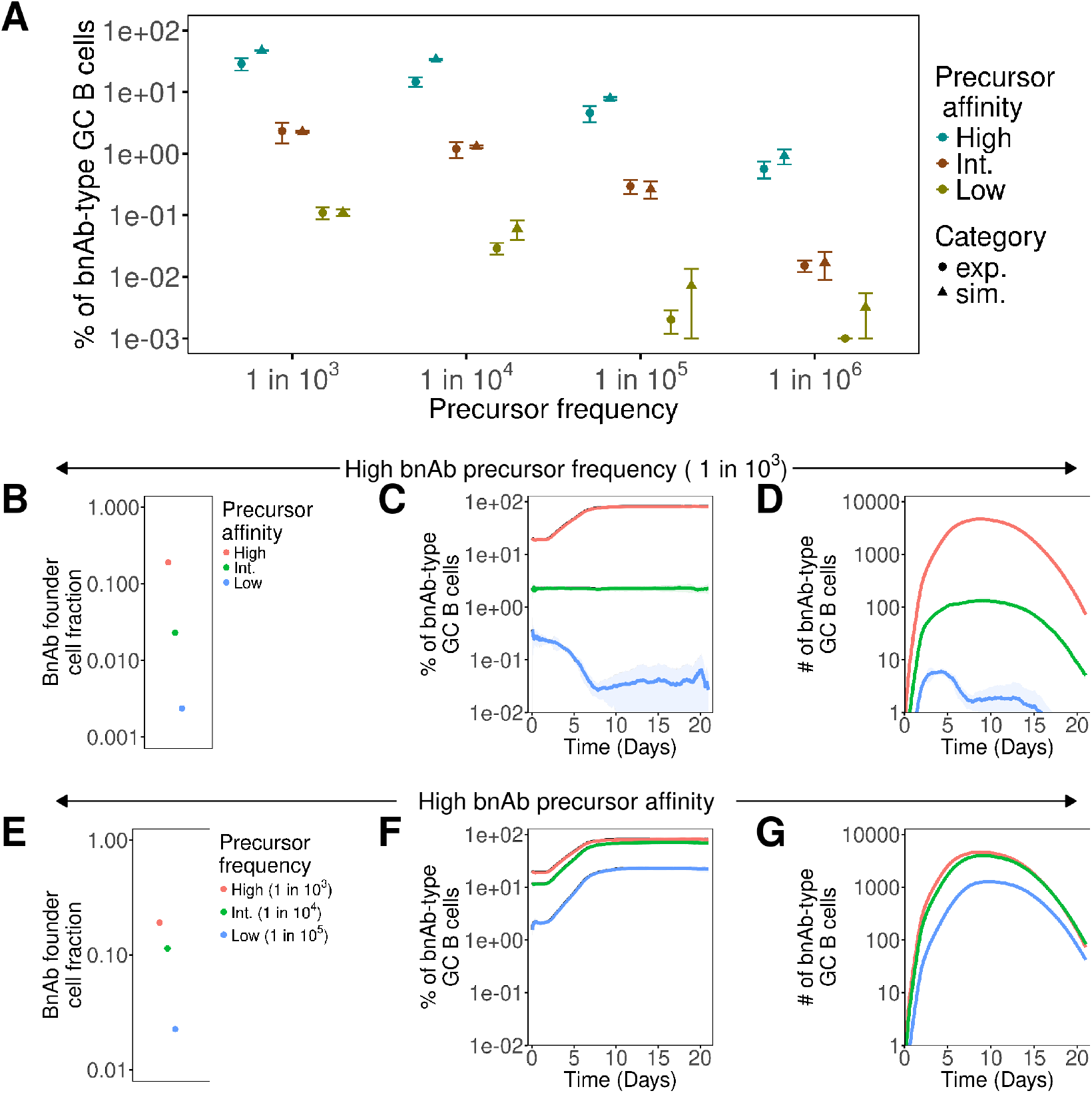
Simulations recapitulate precursor effect as seen in experiments. (A) Comparison of bnAb-type GC B cells in experiments^17^ and simulations. BnAb precursor affinity (*K*_D_) in wildtype C57BL/6 mice was varied as high (0.125*µM*), intermediate (1.3*µM*) or low (18.5*µM*) while frequency was 10-fold serially diluted from 1 in 10^3^ to 1 in 10^6^. Accounting for the 3 days taken for GC onset^95^, simulation data (triangles) at day 5 post GC formation was compared to experimental data (circles) at day 8 post immunization. Extended results for simulations with (B-D) high bnAb precursor frequency (1 in 10^3^) and variable affinity, (E-G) high bnAb precursor affinity and variable frequency. (B, E) Fraction of bnAb-type GC founder cells and temporal curves of (C, F) the percentage and (D, G) number of bnAb-type GC B cells. Means (simulation points and lines) were calculated by averaging the pooled means obtained from 4 sets of 100 simulations. Standard deviation (shaded areas and error bars) was calculated as the standard error of means.

To further elucidate the role of precursor affinity and frequency, we elaborated specific simulation sets from Fig.2A. For the first set, bnAb precursor frequency was high (1 in 10^3^) while the affinity was variable. Under these conditions, the fraction of bnAb-type GC founder cells increased with increasing precursor affinity (Fig.2B). *In silico* GCs seeded by these founder cells showed a similar trend where the percentage and absolute number of bnAb-type GC B cells were more pronounced when precursor affinity was higher (Fig.2C, D). Notably, individual GCs were highly stochastic, with the proportion of bnAb-type GC B cells varying widely(Fig.S2A). A large stochastic variability was also found in experiments, where a singular B cell clone exhibited highly varied degrees of dominance within GCs originating under identical conditions^77,87^. Furthermore, in our simulations, bnAb lineages under high and intermediate precursor affinity conditions affinity matured, in contrast to the low affinity case, where the lineage died out before maturation (Fig.S2B). Evolution of average affinity and total number of all GC B cells were similar in all cases (Fig.S2C, D) and kinetically similar to other independent studies^38,45,88^.

Next, we selected a simulation set where bnAb precursor affinity was high, while the frequency was variable. Here, increased bnAb precursor frequency led to a greater proportion of bnAb-type GC founder and B cells (Fig.2E-G and Fig.S2E). Affinity maturation of bnAb lineages for the considered cases was comparable (Fig.S2F), with minimal differences in the affinity and total number of all GC B cells (Fig.S2G, H). Thus, in our simulations, increasing the relative frequency and affinity of bnAb precursor B cells led to robust priming and improved survival bnAb lineages. This observation was in line with multiple experiments^16–18,22,40,53,89^. Therefore, our model qualitatively captured the precursor effect and reproduced essential GC features.

### Combination of AEM and the precursor effect shapes bnAb lineage response to a multi-epitope antigen

Germline-targeting immunogens such as eOD-GT8 can produce unwanted B cell responses that target epitopes unrelated to those recognized by bnAbs^10,16,19,23,35–37^. We examined priming of bnAb lineages in such a multi-epitope setting next. Here, in addition to the precursor effect, AEM was also expected to modulate GC B cell responses. This modulation was also seen in the earlier single-epitope setting, but to a limited extent. To elucidate this, we repeated simulations with varying precursor affinity performed earlier (Fig.2B-D) without AEM (Fig.S3). While the percentage of bnAb-type GC B cells was unchanged, overall affinity maturation and GC termination were delayed. The latter observations were due to the absence of AEM driven antigen inaccessibility and also noted in a prior study that explored the impact of serum antibodies on GC B cell responses^54^.

Immunogens such as those for HIV can have multiple conformational epitopes that can be classified based on precursor B cells that they bind. Epitopes, such as the CD4 binding site, contain residues that can be contacted by mature bnAbs and their precursor B cells, such as those of the VRC01 bnAb lineage and by non-bnAb precursor B cells as well^10,19,37^. In contrast, epitopes such as the base of the ENV trimer are only contacted by non-bnAb precursor B cells^90–92^. On the eOD-GT8 60mer immunogen, when the highly immunogenic CD4 binding site is blocked, non-bnAb precursor B cells can still bind the antigen, suggesting the presence of epitopes that are not bound by bnAb precursors^10,19,36,37^. To accommodate these features qualitatively, we considered a model antigen (Fig.3A) with two epitopes, referred to as the bnAb and non-bnAb epitopes. The bnAb epitope can bind to bnAb and non-bnAb precursor B cells, while the non-bnAb epitope can only to non-bnAb precursor B cells. Furthermore, we considered the epitopes to be structurally and spatially distinct such that cross-reactivity between B cells and antibodies specific for either epitope could be neglected. These assumptions are reasonable upon consideration of the work by Tas et al.^58^, where post immunization with germline-targeting immunogens, the presence of pre-existing antibodies only disrupted responses of B cells with shared epitope specificity while those specific for alternate epitopes were largely unaffected. Accordingly, model representation of epitopes involved positioning them sufficiently apart on the shape space (see Antigen-presentation by FDCs). Moreover, we assumed frequencies and affinities of non-bnAb precursor B cells for both epitopes to be equal, as related data for wildtype C57BL/6 was unavailable, although this assumption was relaxed in later sections. We recognize that T cell signaling may also bias B cell responses towards certain epitopes^93,94^. However, as our focus was on the precursor effect and AEM, we assumed equal T cell help for B cells irrespective of epitope specificity.

**Figure 3.**
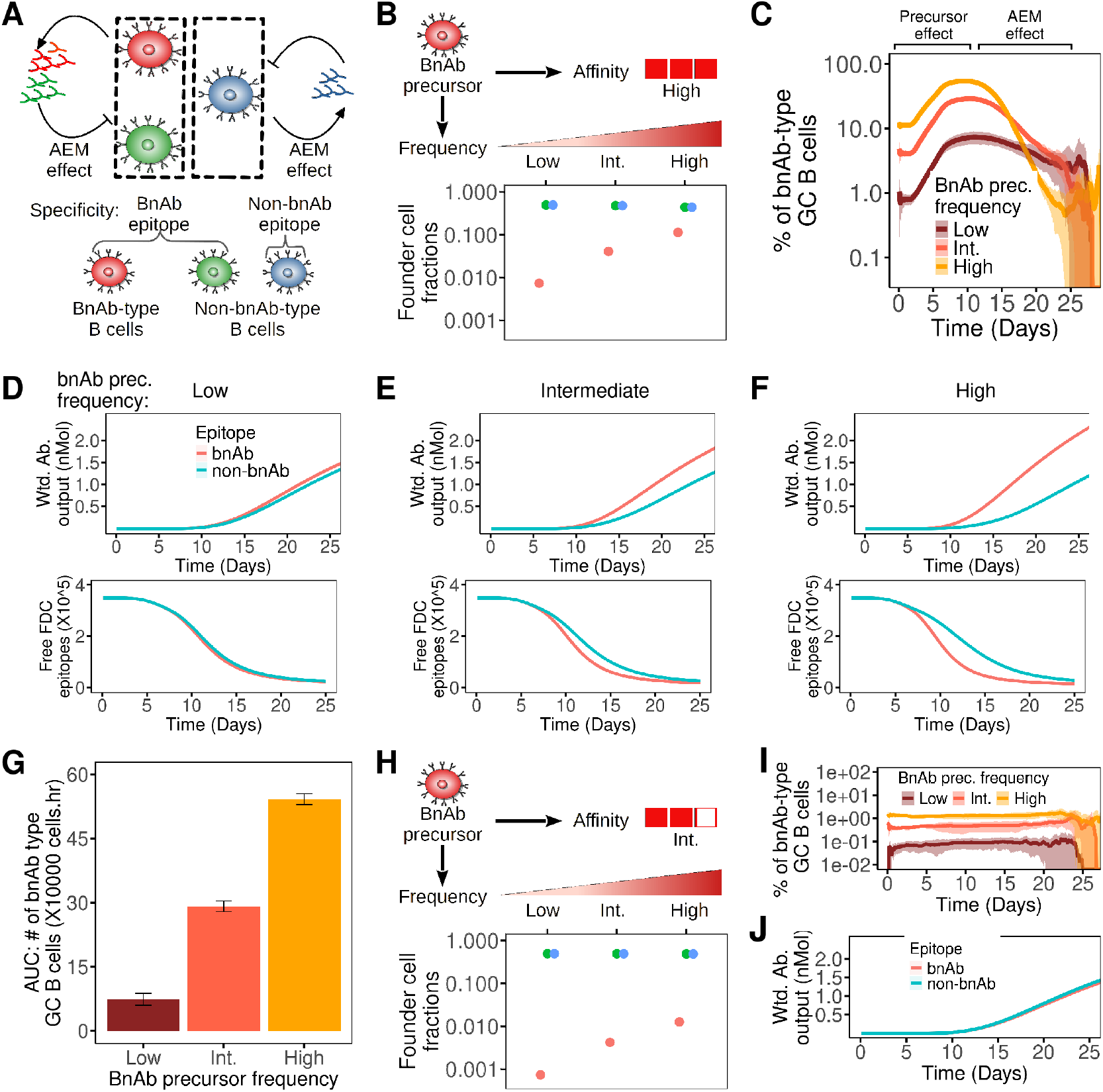
AEM modulates B cell responses in multi-epitope settings –. (A) Simulation schematic with epitope-specific AEM. BnAb precursor frequency was varied as high (1 in 10^4^), intermediate (2 in 10^5^) and low (3 in 10^6^) while affinity (*K*_D_) was fixed at (B-G) high (0.400*µM*) or (H-J) intermediate (1.80*µM*). GC characteristics: (B) founder cell fractions, (C) percentage of bnAb-type B cells, (D-F (top)) endogenous antibody output weighted by epitope-specific affinity and (D-F (bottom)) available free antigenic epitopes on FDCs for (D) low, (E) intermediate and (F) high bnAb precursor frequencies, (G) AUC of bnAb-type B cells. GC characteristics for intermediate bnAb precursor affinity (1.80*µM*) simulations: (H) founder cell fractions, (I) percentage of bnAb-type B cells and (J) endogenous antibody output weighted by epitope-specific affinity. Means (bars and lines) were calculated by averaging the pooled means obtained from 4 sets of 100 simulations. Standard deviation (shaded area and error bars) was calculated as the standard error of means.

Accordingly, we evaluated GC B cell responses by considering the wildtype C57BL/6 mouse system, where affinity and frequency of bnAb precursor B cells were modulated. To ensure consistency with data from Huang et al.^17^, simulations in (Fig.2) were repeated with the two epitope model. The model still recapitulated the experimental data (Fig.S4), indicating its overall robustness.

Next, the bnAb precursor affinity was set high (0.400*µM*) while the frequency was varied as high (1 in 10^4^), intermediate (2 in 10^5^) and low (3 in 10^6^). Based on this, founder cell fractions were calculated (Fig.3B) and used to simulate the GC reaction. Due to the precursor effect, increasing precursor frequency improved the percentage of bnAb-type GC B cells for a substantial part of the GC response (Fig.3C). Strikingly however, this percentage declined in the later GC phase due to AEM. This was in contrast to earlier single-epitope simulations (Fig.2C, F) where analogous responses were mostly monotonic after ∼ day 8. When the fraction of bnAb-type GC B cells was large, i.e., for high or intermediate precursor frequency, the antibody pool generated by GC-derived plasma cells became skewed towards the bnAb epitope (Fig.3D-F). This resulted in preferential masking of this epitope, causing its availability in the GC to decline faster. Consequently, the bnAb lineage was inhibited. As an estimate of the cumulative response, the AUC (area under curve) based on absolute numbers of bnAb-type GC B cells was calculated over the entire response duration using hourly values. The AUC was much larger at higher precursor frequencies (Fig.3G), indicating that gains due to the precursor effect superseded AEM-mediated inhibition.

We also simulated with bnAb precursor affinity as intermediate (1.80*µM*) and varied the frequency as before (Fig.3H). The initial percentage of bnAb-type GC B cells was now about an order of magnitude lower in comparison to the high precursor affinity setting due to the precursor effect (Fig.3I). As a consequence, antibodies were minimally biased for the bnAb epitope even when the precursor frequency was high (Fig.3J), resulting in weak AEM.

To check whether these signatures of AEM-mediated regulation were consistent with experiments, we considered the study by Wang et al.^18^, where wildtype C57BL/6 mice with bnAb precursor frequencies and affinities similar to the values tested in Fig.3 were immunized with eOD-GT8 60mer. In agreement with our simulation results, when bnAb precursor frequency and affinity were high in experiments, the fraction of bnAb-type GC B cells at day 36 was lower in comparison to the fraction at day 8 post immunization. More precisely, these differences were statistically significant at the 5% significance level, as reflected by their p values of 0.00221 and 0.00123 for experiments performed with high affinity precursor B cells CLK21 (0.440*µM*) and CLK09 (0.350*µM*), when their frequencies were high (1 in 10^4^) and intermediate (2 in 10^5^) Table1. Importantly, when precursor frequency and/or affinity were reduced, differences in late to early fractions of bnAb-type GC B cells were not statistically significant. For further validation, we performed a corresponding analysis on the simulation data. As in silico GCs rapidly extinguished around day 25, day 24 and day 5 post GC formation were chosen as being representative of late and early GCs, accounting for 3 days needed for GC formation^95^. The pattern and the significance of p-values were comparable in experiments and simulations.

**Table 1:**
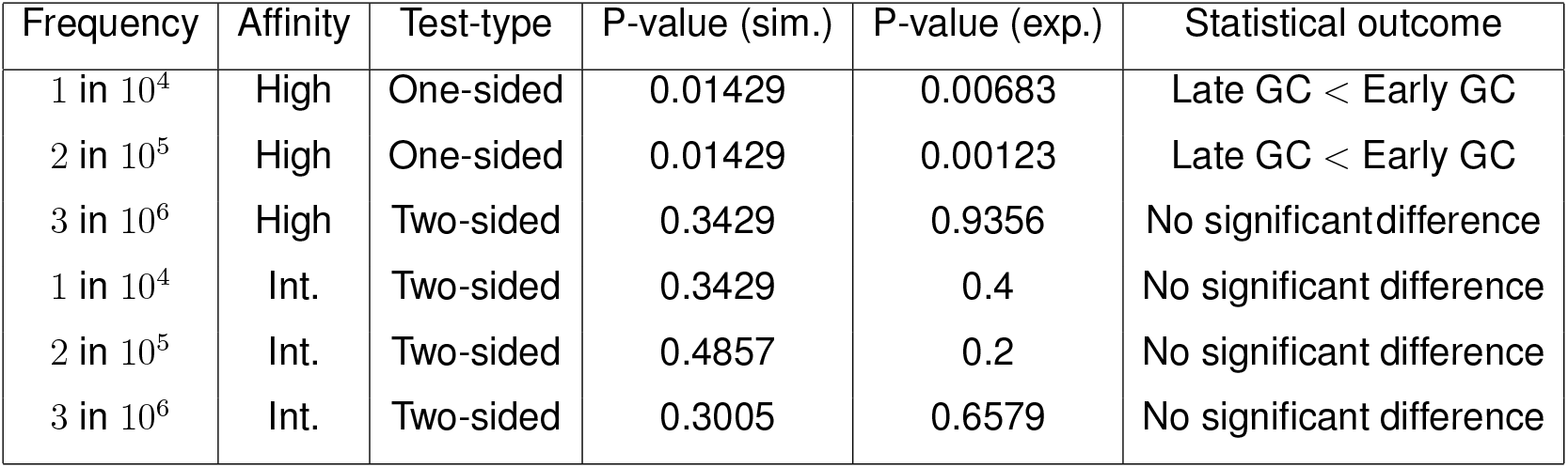
P-values obtained from non-parametric Wilcoxon Rank-Sum (Mann-Whitney U) test. BnAb precursor frequencies and affinities were varied and the resultant percentages of bnAb-type GC B cells were compared at late vs. early GC time points for experiments^18^ (Fig.6 B of reference) and earlier simulations in Fig.3. For high affinity scenarios, experimental data of CLK21 (.440*µM*) and CLK09 (.350*µM*) precursors was combined. Time points chosen were day 8 vs. day 36 post immunization for experiments and day 5 vs. day 24 post GC onset for simulations (for further details, see text). The one-sided test was used to assess a specific directional difference between early and late GC responses, while the two-sided test was used to evaluate any statistically significant difference regardless of direction. The tests were performed using the wilcox.test function in R.

Thus, the above results provide important evidence that AEM modulates evolution of bnAb lineages in addition to the precursor effect. Overall, our model qualitatively reproduced the trends of bnAb-type GC B cell kinetics as seen in experiments employing physiologically relevant preclinical mouse models, indicating its utility and robustness^17,18^. In this context, an important advantage of our model is that undesirable non-bnAb-type GC B cells can also be tracked. We examined their evolution next.

### AEM contributes to resilience of B cell responses to non-bnAb epitopes

To assess responses of non-bnAb-type GC B cells we re-considered the simulations performed with high bnAb precursor affinity and variable frequency (Fig.3B-G). Due to the precursor effect, we expected that simulations with increased bnAb precursor frequency would exhibit reduced responses of non-bnAb-type GC B cells. Our calculations confirmed this expectation as the percentage of non-bnAb-type GC B cells (Fig.4A-C) and associated AUCs (Fig.4D, E) declined when bnAb precursor B cells were more abundant. However, the AUC of non-bnAb-type GC B cells that were specific for the bnAb epitope declined much more, about ∼ 53%, compared to a decline of only ∼ 17% for GC B cells of non-bnAb epitope specificity. We attributed these observations to AEM: for high precursor frequency settings, as the endogenous antibody pool was skewed towards the bnAb epitope, its availability and cognate B cell population were diminished. However, such regulation was minimal for non-bnAb-type GC B cells specific for the non-bnAb epitope.

**Figure 4.**
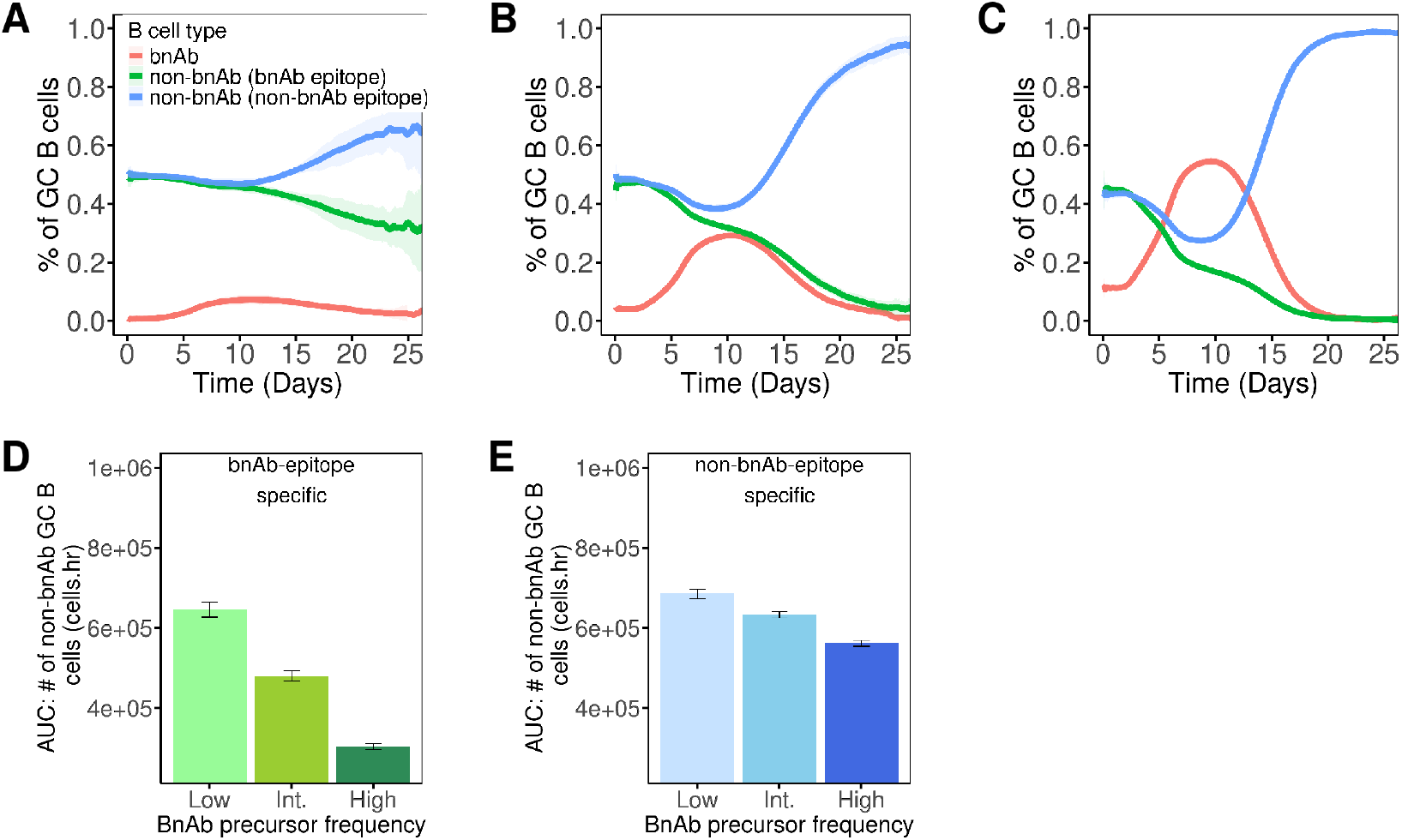
Evolution of non-bnAb-type GC B cell responses in relation to Fig.3A-G. BnAb precursor frequency was varied as (A) low, (B) intermediate and (C) high while affinity was fixed at high (0.400*µM*). Temporal trends of GCs as: (A-C) percentage of bnAb- and non-bnAb-type GC B cells, AUC of non-bnAb-type GC B cells specific for (D) bnAb epitope and (E) non-bnAb epitope. Means (bars and lines) were calculated by averaging the pooled means obtained from 4 sets of 100 simulations. Standard deviation (shaded area and error bars) was calculated as the standard error of means.

These results indicate that potentiating bnAb precursor B cells caused B cell competition to manifest differently for non-bnAb-type GC B cells depending on their epitope specificities. Motivated by these observations, we asked the inverse question, i.e., can epitope-specific non-bnAb B cells exert different competitive pressures on the bnAb lineage? Our model is well suited to address this as it can accommodate epitope-specific changes in properties of non-bnAb precursor B cells. We present such an analysis next.

### Non-bnAb precursor B cells inhibit bnAb lineage output, with stronger inhibition by cells of shared epitope specificity

In all two epitope simulations so far, we assumed equal epitope-specific frequencies of non-bnAb precursor B cells. However, these frequencies are likely different for different immunogens.

To predict the impact of these variations, we considered two extreme scenarios. For the first scenario, we assumed non-bnAb precursor B cells to be biased for the bnAb epitope (90%), while for the second scenario, the same bias was assumed for the non-bnAb epitope (Fig.5A). To mimic human physiology, precursor features of the bnAb lineage were selected based on those of human VRC01 bnAb precursor B cells. We only considered precursor B cells having substantial affinity (*K*_D_ *<* 3*µM*)^9,10^ as associated responses are most relevant for GCs^16–18^. This translated to a low bnAb precursor frequency (∼ 1 in 10^6^) and a harmonic mean precursor affinity of 0.195*µM* (harmonic averaging is done due to the inverse relationship of affinity with *K*_D_, see Pre-GC Model). All other systemic and antigenic properties were consistent with the earlier two epitope simulations.

**Figure 5.**
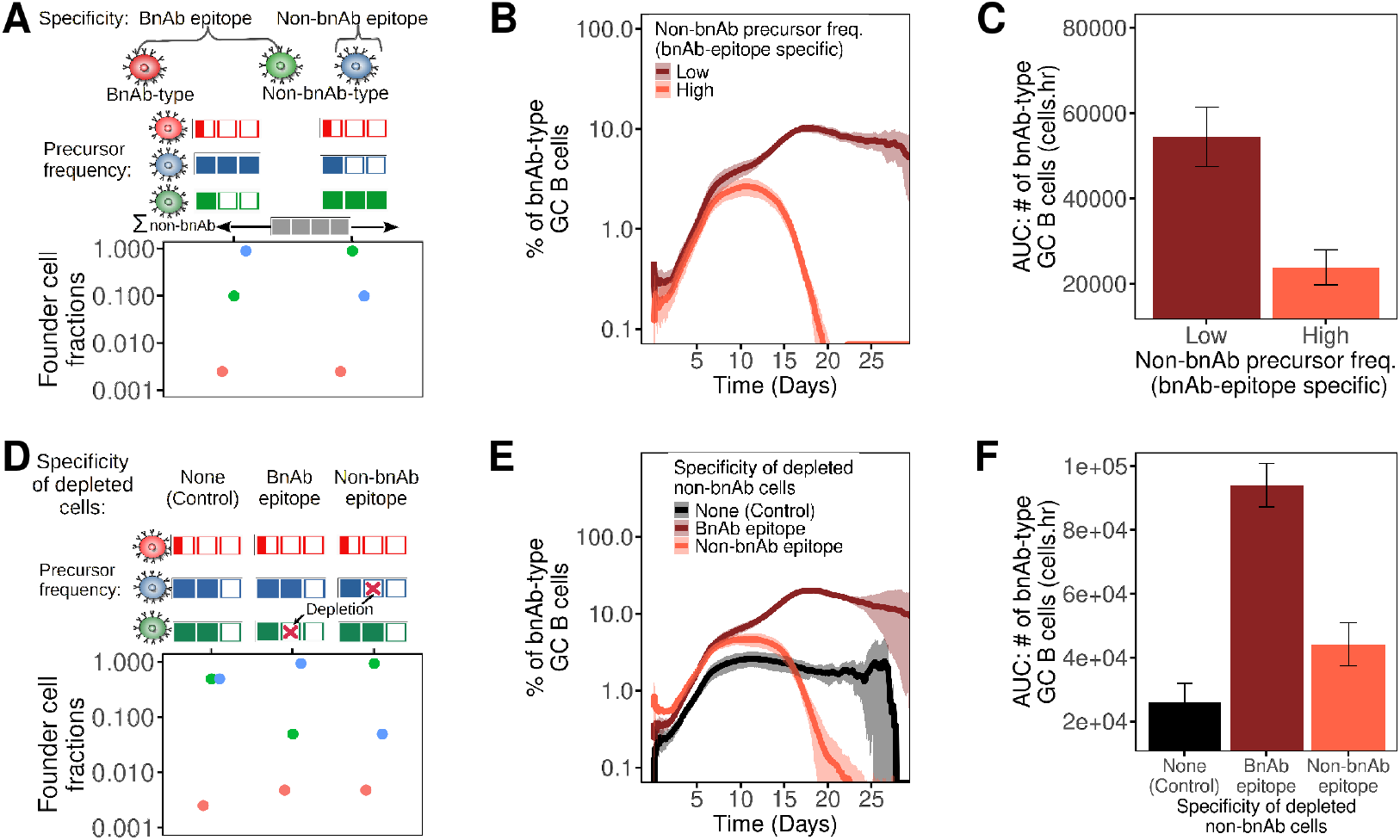
Impact of non-bnAb precursor B cells on bnAb lineage priming. (A-F) BnAb precursor frequency was low (1 in 10^6^) and harmonic mean affinity was high (0.195*µM*). (A-C) 10% (low) or 90% (high) non-bnAb precursor B cells were specific for the bnAb epitope. (D-F) Non-bnAb-type GC founder cells were depleted by 90% in an epitope-specific manner (control case: equal targeting of both epitopes by non-bnAb-type GC B cells). GC B cell characteristics: (A, D) founder cell fractions, (B, E) percentage of bnAb-type GC B cells and (C, F) AUC of bnAb-type GC B cells. Colored boxes represent classes of the frequencies of non-bnAb precursor B cells and do not reflect actual values or scale. Means (bars and lines) were calculated by averaging the pooled means obtained from 4 sets of 100 simulations. Standard deviation (shaded area and error bars) was calculated as the standard error of means.

BnAb-type GC B cells evolved similarly for both scenarios until ∼ day 12 (Fig.5B). This was because the precursor effect impacted both scenarios equally, a consequence of the overall affinity and frequency of bnAb and non-bnAb precursor B cells being identical. Strikingly however, in the late GC phase, the percentage of bnAb-type GC B cells diverged and was lower when non-bnAb precursor B cells specific for the bnAb epitope were dominant. This observation was a consequence of AEM: GC B cells and related endogenous antibodies were biased towards the bnAb epitope, resulting in its masking and the consequent reduction of bnAb-type GC B cells (Fig.5B, C).

In the above calculations, the total frequencies of bnAb and non-bnAb precursor B cells were unchanged. However, it is possible to modify immunogens such that binding of non-bnAb precursor B cells is inhibited^96,97^. By implementing strategies such as glycan masking, variants of eOD-GT8 with ∼ 90% reduction in B cell responses towards alternate non-bnAb epitopes have been developed^36^. Here, we evaluated whether such a selective depletion of non-bnAb precursor binding improves bnAb-type GC B cell responses.

To model an immunogen with diminished binding of non-bnAb precursor B cells, we first assumed their precursor frequencies (and consequently their founder cell fractions) for both epitopes to be equal. Subsequently, non-bnAb founder cell fractions were reduced by 90%, focusing on one epitope at a time. BnAb precursor frequency and affinity were kept unchanged.

Simulations performed with these conditions, showed that bnAb-type GC B cells became more dominant when binding of non-bnAb precursor B cells was reduced (Fig.5D-F). This gain was highest when antigen binding of non-bnAb precursor B cells that are bnAb epitope-specific was compromised, as AEM directed towards the bnAb epitope was also reduced. Thus, our simulations predict that at physiologically relevant bnAb precursor frequencies and affinities, immunogens with reduced binding of non-bnAb precursor B cells are expected to make bnAb-type GC B cells more dominant due to the precursor effect. Importantly, these GC responses are further improved when binding of non-bnAb precursor B cells specific for the bnAb epitope is diminished, as here, AEM is also supportive.

The above analysis demonstrates the utility of our model in predicting bnAb lineage evolution under various conditions and identifying those that are likely to yield improved responses. As single bnAb lineages are unlikely to be sufficient for protection against HIV^30,31,31–33^, we next explored scenarios with multiple lineages.

### At low precursor frequencies, simultaneous priming of multiple bnAb lineages is additive

Priming of numerous bnAb lineages, defined in this work as the activation and subsequent maturation of their precursor B cells in an ensemble of GCs, may be achieved by either (i) a single immunogen targeting precursor B cells of multiple lineages simultaneously or (ii) by employing multiple immunogens, each targeting precursor B cells of an individual lineage. Here, we evaluated both possibilities while varying the features (affinity, frequency and epitope-specificity) of bnAb and non-bnAb precursor B cells.

We considered a model antigen system with two epitopes (epitope 1 and epitope 2), each of which may bind to bnAb and non-bnAb precursor B cells (Fig.6A). For scenario (i), multiple bnAb lineages were primed simultaneously with a single immunogen by simulating concurrent maturation of lineages in the same GC ensemble. In contrast, for scenario (ii), to simulate priming with distinct immunogens, each bnAb lineage was stimulated individually (maturation in different GC ensembles) and the resultant bnAb-type responses were summed to get the aggregated output. In line with recent works^98,99^, such a formulation assumes that interference between responses to different immunogens is minimal and thus predicts outputs that are in the upper limit of what may be realized under practical constraints of immunization. For a meaningful comparison, identical sets of non-bnAb precursor B cells were assumed in both scenarios. All other fundamental assumptions were consistent with previous two epitope simulations.

**Figure 6.**
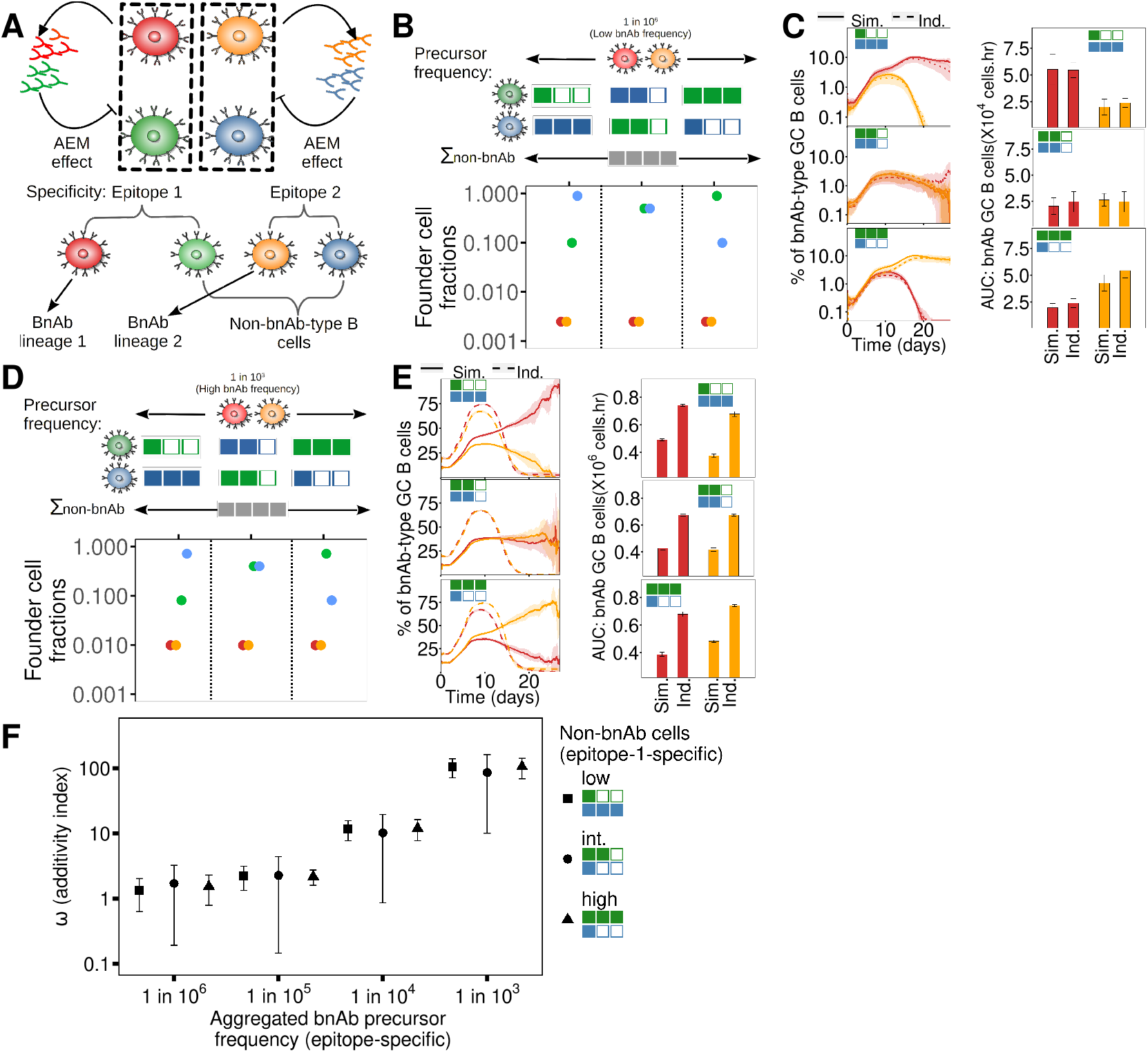
Simultaneous vs. individual priming of multiple bnAb lineages with distinct epitope specificities. (A) Simulation schematic for multiple lineages with AEM. For simultaneous priming, bnAb lineages are simulated concurrently in a single GC ensemble, while for individual priming, each GC ensemble contains a singular bnAb lineage. (B-F) 90%, 50% or 10% non-bnAb precursor B cells were specific for epitope 1. For both bnAb lineage 1 (red) and 2 (orange), harmonic mean precursor affinity was high (0.195*µM*) while frequency was low (B, C, 1 in 10^6^) or high (D, E, 1 in 10^3^). GC B cell characteristics: (B, D) founder cell fractions, (C, E left) percentage of bnAb-type cells for simultaneous (solid lines) or individual (dashed lines) lineage priming and (C, E right) AUC of bnAb-type cells. (F) Ω (additivity index) calculated for an arbitrary number of bnAb lineages. Colored boxes represent classes of the frequencies of non-bnAb precursor B cells and do not reflect actual values or scale. Means (bars and lines) were calculated by averaging the pooled means obtained from 4 sets of 100 simulations. Standard deviation (shaded area and error bars) was calculated as the standard error of means.

We first evaluated simultaneous priming with a single immunogen (scenario (i)) of two bnAb B cell lineages that were annotated as lineage 1 or 2 based on their epitope specificity. Precursor features of both lineages were based on the precursor B cells of the VRC01 bnAb having substantial affinity (*<* 3*µM*) in humans (frequency: 1 in 10^6^ and harmonic mean affinity: 0.195*µM* ^9,10^). Furthermore, epitope specificities of non-bnAb precursor B cells were varied, i.e., 90%, 50% or 10% of these precursors were biased towards one of the two epitopes, with the rest being specific for the alternate epitope.

Simulation results showed that due to manifestation of the precursor effect, initial trajectories of both lineages were similar for all cases (Fig.6B, C). However, AEM, driven by differential epitope specificities of non-bnAb precursor B cells, modulated bnAb-type responses in late GCs. When non-bnAb precursor B cells were biased for epitope 1, response of lineage 1 declined while lineage 2 was more pronounced. When the bias of non-bnAb precursor B cells was changed to favor epitope 2, these observations were reversed.

Next, we performed analogous simulations for scenario (ii), where bnAb lineages were primed individually with distinct immunogens (maturation in different GC ensembles). The impact of non-bnAb precursor B cells in these simulations was similar to earlier (scenario (i)) simulations. Importantly, responses of bnAb lineages were comparable in both scenarios, indicating that simultaneous priming of bnAb lineages (concurrent maturation in the same GC ensemble) did not compromise lineage output as compared to individual priming (maturation in different GC ensembles) for the tested conditions.

In an extreme scenario, we analyzed bnAb lineages with supraphysiological precursor frequency (1 in 10^3^) while keeping all other settings and analysis methodology the same (Fig.6D, E). The modulation of bnAb-type GC B cells by AEM was similar but dampened in comparison to the earlier low frequency simulations. Moreover, in contrast to the earlier simulations, simultaneous priming (scenario (i)) yielded lower bnAb-type GC B cell responses when evaluated against individual priming (scenario (ii)).

As germline-targeting strategies can be potentially tailored to target precursor B cells of multiple distinct bnAbs, we next evaluated a more generalized setting involving an arbitrary number of bnAb lineages. Simulating this requires bnAb lineage-specific precursor frequencies, which are often unquantified for humans. To circumvent this, for simultaneous priming (scenario (i)), simulations were performed based on aggregated epitope-specific bnAb precursor frequencies in a single GC ensemble (for detailed description see (Simulation formulation for arbitrary number of bnAb lineages)). To simulate scenario (ii), these aggregated frequencies were split into smaller precursor subsets that were defined ubiquitously in the lower frequency limit (1 in 10^6^), with each subset specific for a single epitope. Outputs from simulations of these individual subsets in different GC ensembles were then summed to obtain the AUC. The aggregation and splitting of bnAb precursor B cells is reasonable as only high affinity bnAb precursor B cells (harmonic mean affinity: 0.195*µM*) were considered. Also, the overall epitope-specific split of bnAb precursor B cells was assumed to be equal. This simulation setup thus represents the theoretical extremes of simultaneous and individual bnAb lineage priming.

For comparison, the additivity index Ω was defined as the ratio of AUCs obtained from simultaneous priming versus individual priming. For Ω ≈ 1, lineage responses were “additive” meaning simultaneous priming generated responses comparable to the sum of responses obtained when subsets of bnAb precursor B cells were individually targeted. In contrast, Ω *>* 1 meant lineage responses were comparatively suppressed when primed simultaneously. With this formulation, simulations performed with aggregated epitope-specific bnAb precursor frequency varied from physiologically relevant (1 in 10^6^) to supraphysiological (1 in 10^3^) values yielded Ω close to 1 for lower (*<* 1 in 10^5^) precursor frequencies. For higher frequencies, Ω increased sharply (Fig.6F). As a consistency check, all the above results were valid even when bnAb precursor B cells were specific only for epitope 1 (Fig.S5).

Overall, these results indicate that priming of multiple bnAb lineages is modulated by the precursor effect and AEM. Additionally, at physiological, i.e. low, precursor frequencies, bnAb-type responses are additive, meaning individual lineages are not diminished when targeted simultaneously with a single immunogen.

## DISCUSSION

By limiting the total number of distinct proteins needed, individual immunogens capable of adequately priming precursor B cells of multiple bnAbs would be an important advance for HIV vaccinology. In this work, we evaluated whether intrinsic mechanisms of B cell competition and selection can compromise responses generated by such complex immunogens. Using an *in silico* simulation framework of pre-GC and GC phases, we showed that the priming of bnAb lineages is impacted by (i) features of antigen binding precursor B cells, i.e., frequency and affinity (precursor effect) and (ii) masking of cognate epitopes by antibodies derived from GC-exiting plasma cells (AEM). Due to the precursor effect, lineages with increased precursor frequency and/or affinity were more competitive and therefore exhibited improved cumulative GC responses. In contrast, when B cell responses were initially biased to the bnAb epitope, AEM obscured the epitope and inhibited cognate B cell responses, including those of the bnAb lineage in the later GC phase. Based on this, our simulation framework qualitatively recapitulated features of bnAb lineage evolution as observed in studies using pre-clinical mouse models with the eOD-GT8 60mer immunogen^17,18^. Importantly, in simulations performed with physiologically comparable bnAb precursor frequencies (*<* 1 in 10^5^), multiple bnAb lineages could be primed simultaneously with a single immunogen in the same GCs without loss of individual lineage output.

For a protective HIV vaccine, high neutralization potency and titres of antibody responses would be required^3,26,30^. Adequate priming of bnAb precursor B cells followed by boosting with appropriate immunogens will be instrumental in achieving this^7,8,23,24,26–29^. However, diversion of B cell responses towards non-bnAb-type cells can be disruptive^23,26,35,36^. In our simulations, when the bnAb lineage was potentiated due to increased precursor frequency and affinity (Fig.3), the corresponding GC response improved because undesirable non-bnAb-type GC B cells were inhibited due to the precursor effect. However, this led to AEM being directed towards the bnAb epitope, resulting in inhibition of epitope-specific bnAb- and non-bnAb-type GC B cells in the later GC phase. Moreover, as AEM-mediated inhibition of the non-bnAb epitope was relatively low, its cognate B cells were much more resilient (Fig.4). Indeed, this resilience of B cell responses to non-bnAb epitopes after a single immunization with eOD-GT8 60mer is noted in multiple mouse models that have high frequencies of bnAb precursor B cells^23,35,36^. Our simulations here provide a mechanistic basis for these observations via AEM.

Furthermore, in our simulations, diminishing the antigen-binding non-bnAb precursor B cells (Fig.5D-F) generally improved bnAb-type GC B cell responses due to the precursor effect. Therefore, we predict that further design modifications in germline-targeting immunogens to eliminate or hide non-bnAb epitopes, such as in^36^, should improve bnAb lineage priming at physiologically relevant precursor frequencies and affinities.

The simulation results suggested that AEM-mediated inhibition of the bnAb epitope and its cognate B cells could occur when non-bnAb precursor B cells were largely (∼ 90%) specific for it (Fig.5). Accordingly, in simulations where the proportion of bnAb epitope-specific non-bnAb precursor B cells was reduced or when these cells were specifically eliminated, AEM was mitigated and bnAb-type GC B cell responses were optimized. These results are also relevant clinically as ∼ 90% eOD-GT8-binding naive B cells are specific for the bnAb epitope (CD4 binding site) in humans^10^. Thus, we expect substantial improvements in bnAb-type GC B cell responses if eOD-GT8 is modified to reduce binding of non-bnAb precursor B cells that are bnAb epitope-specific. In order to evaluate whether AEM-mediated suppression is relevant for other immunogens, including those that target multiple lineages such as GT1, estimation of epitope specificities of antigen-binding non-bnAb precursor B cells would be needed. If these cells are biased for bnAb epitopes, AEM would be relevant and bnAb lineage priming is expected to be suboptimal.

In previous works, pre-existing antibodies sourced from primary immunization or external injections have been implicated in increasing antibody breadth by diversifying B cell responses towards alternate poorly targeted antigenic epitopes^55,57–61,67,100–102^. Our findings here extend these observations by demonstrating that even primary GCs can be modulated by contemporary endogenous antibodies via AEM. They also suggest that by distracting B cell responses towards alternate non-bnAb epitopes, AEM can be counter-productive to germline-targeting strategies, as these strategies usually involve modification of the antigen such that the immunogenicity of the bnAb epitope is enhanced^13,36,37,97^.

In addition to high titres, combinations of bnAbs that can neutralize multiple HIV strains would also be required for a protective vaccine^31,32,103^. BnAbs that target distinct ENV epitopes show complementarity in neutralization and thus can be instrumental in achieving this^33^. The GT1 immunogen and associated variants that are based on intact ENV trimers are important in this context as they can target bnAb precursor B cells that are specific for the CD4 binding site and the V2 apex^34^. Through further modifications, it might be possible to target additional epitopes as was demonstrated for the recently developed 3nv.2 immunogen which targets bnAb precursors for three distinct epitopes^104^. Additionally, targeting precursor B cells of bnAb lineages with shared epitope specificity may also be advantageous, as different lineages can have different requirements of SHM and rare mutations^3,5^. For example, the IOMA class and the VRC01 class bnAbs are both specific for the CD4 binding site and their precursor B cells can bind to eOD-GT8 60mer^10^. As IOMA class bnAbs require fewer insertions, deletions and overall mutations, compared to the VRC01 bnAb, they could have an easier evolutionary trajectory from a boosting perspective^5,37,105^. However, VRC01 has larger breadth and potency, a quality that is also shared by other bnAbs having extensive SHM and rare mutations^5^. Overall, an immunogen capable of priming multiple lineages would be compatible with many possible paths for bnAb induction.

In this context, our results indicate that at physiologically relevant low bnAb precursor frequencies (*<* 1 in 10^5^)^9–11^ overall bnAb-type GC B cell responses were similar regardless of whether the lineages were primed simultaneously with a single immunogen or individually with distinct immunogens (Fig.6 and Fig.S5). However, this additivity ceased at higher frequencies, where simultaneous priming generated comparatively inferior responses. We note here that, since our model was calibrated using data from experiments in mice, the precise frequency at which simultaneous priming becomes inferior is expected to change in humans depending on the precursor pool targeted by immunogens. However, as a general principle, simultaneous priming was efficient at low precursor frequencies across variations in frequencies, affinities and epitope specificities of the sets of antigen-binding precursor B cells. Thus, these results make a case for the further development of immunogens that activate multiple precursor B cells belonging to different bnAb lineages.

### Limitations of the study

By integrating the precursor effect and AEM, our model framework provides fundamental insights into the mechanics of bnAb lineage responses while predicting scenarios where optimal priming of individual and multiple lineages can be achieved. Although this work focused on the precursor effect and AEM, B cell responses are also modulated by antigen avidity^53,106^. Incorporation of the avidity effect would be relevant for predicting responses generated by lower avidity immunogens. Furthermore, we assume that all participating B cells have identical probabilities of acquiring mutations and associated affinity changes. Implementing bnAb-lineage-specific mutation and affinity landscapes may improve prediction accuracy for specific lineages and allow tracking of B cell populations that acquire rare but important bnAb-like mutations^20,94^. Lastly, we did not investigate the role of additional variables that shape GC B cell competition and selection, such as timing, duration and dose of antigen delivery^90,107,108^ and epitope-specific biases of Tfh cells^93,94^. Although the insights of this study are sufficiently general, integration of these variables into the present model can help in isolating conditions that further optimize bnAb lineage priming.

## Supplemental information index

Figures S1-S5 and their legends in a PDF

## Acknowledgments

AKG was supported by the Innovative Medicines Initiative 2 Joint Undertaking (JU) under grant agreement No 101007799. The JU receives support from the European Union’s Horizon 2020 research and innovation programme and EFPIA. The funding bodies had no role in the design of the study, collection, analysis, and interpretation of the results, or writing the manuscript. The authors thank Dr. Gustavo Hernandez-Mejia for critically reviewing the manuscript and all members of the SIMM group for helpful suggestions.

## Author contributions

Conceptualization, A.K.G., S.C.B., and M.M.H.; methodology, A.K.G., S.C.B., and M.M.H.; investigation, A.K.G.; writing – original draft, A.K.G.; writing – review & editing, A.K.G. and S.C.B. and M.M.H.; funding acquisition, S.C.B. and M.M.H.; resources, S.C.B. and M.M.H.; supervision, S.C.B. and M.M.H.

## STAR METHODS

### Key resources table

*To create the KRT, please use the KRT webform or the Word template and upload this file separately*.

### Resource availability

#### Lead contact

Further information and requests can be addressed to Michael Meyer-Hermann (mmh@theoretica biology.de).

#### Materials availability

This study did not generate new materials.

#### Data and code availability

- This paper analyzes existing, publicly available data.References to the datasets are found in the respective figure legends and are listed again in the key resources table.
- All original code has been deposited at Zenodo and is publicly available as of the date of publication. DOIs are listed in the key resources table.
- Any additional information required to reanalyze the data reported in this paper is available from the lead contact upon request.

### Experimental model and study participant details

This study did not involve any experiments and exclusively relies on computer simulations.

### Method details

We present here the detailed description of the mathematical model used to simulate pregerminal centre (GC) and GC phases of the B cell response.

#### Pre-GC Model

Post immunization, precursor B cells residing in secondary lymphoid organs compete to acquire antigen and CD4 T cell signals, following which they can get activated and seed GCs as founder cells. In this pre-GC phase, as precursors that are more populous or have higher affinity are more likely to get these signals^40,50,51,53^, we assumed activation of these cells to be proportional to their abundance and affinity and formulated relevant equations to calculate the composition of GC-seeding founder cells. The abundance of precursor B cells is commonly expressed as their frequency, which is a ratio of the cell population of interest relative to the overall naive B cell repertoire. Thus, for a scenario consisting of *n*_L_ bnAb lineages and non-bnAb precursor B cells that can bind antigen, the adjusted precursor fraction of the *i*^th^ bnAb lineage, 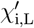, is

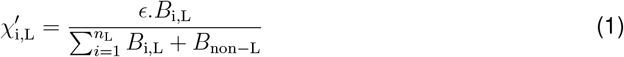

where *B*_i,L_ and *B*_non−L_ are frequencies of antigen-specific precursor B cells of the *i*^th^ bnAb lineage and of non-bnAb-type respectively. In line with the observed murine precursor frequencies for many common antigens^37^, *B*_non−L_ was assumed to be 1 in 10^4^. The error factor, *ϵ*, is used to calibrate the model and is constrained between [0, 1]. It is calculated using experimental data at the end of this section. As a consequence of Equation 1, the adjusted fraction of non-bnAb precursor B cells, 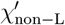, was calculated as 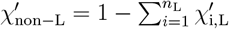

B cell activation and migration to the GC compartment are affinity dependent (inverse dependence on the equilibrium dissociation constant (*K*_D_))^40,50^. Accordingly, the final fraction of activated bnAb-type founder cells, 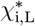, was computed by scaling down adjusted precursor fractions by their respective *K*_D_ values as

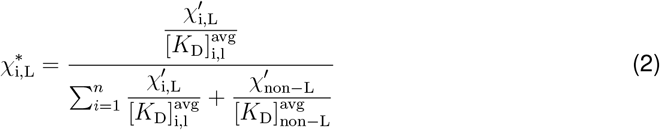

where 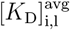 and 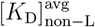 are harmonic means of *K*_D_ values of precursors of the *i*^th^ bnAb lineage and non-bnAb-type respectively. The final Equation 3 that is used to calculate founder cells is obtained by rearranging terms in Equation 2

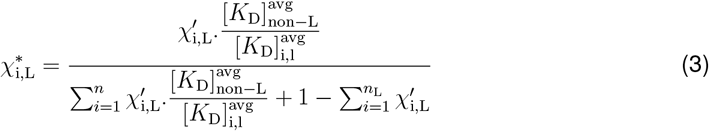

In this work, we mainly model responses in wildtype C57BL/6 mice. The affinities of non-bnAb precursor B cells that bind to germline-targeting immunogens in these mice could not be found in literature. Therefore, to estimate the average affinity, 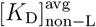, we assumed murine non-bnAb precursor B cells to be distributed uniformly in the *K*_D_ = 10*µM* to 1*µM* affinity range. This range is similar for human non-bnAb precursor B cells that are specific for eOD-GT8, although cells of much lower affinities were also noted^10,19^. However, in our analysis, we ignore possible cells at the lower end of the affinity spectrum as they are expected to be insignificant for GCs at physiologically relevant frequencies^16,17^. Accordingly, 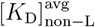 was calculated as being equivalent to 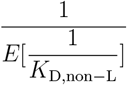 where *K*_D,non−L_ is the affinity of individual non-bnAb precursor B cells. For uniform distribution between 10 and 1*µM*, 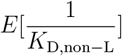 evaluates to 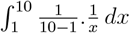 which equates to 0.256. Accordingly, 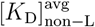 is estimated as ∼ 3.9*µM*.

As bnAb lineages in Fig.2, 3 and 4 are seeded by a single precursor-type, 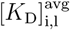 is equal to the bnAb precursor’s affinity. For simulations in Fig.5 and 6, 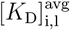 for high affinity bnAb lineage is estimated using precursor affinities of the VRC01 bnAb that are obtained using the eOD-GT8 immunogen in humans^9,10^. Specifically, precursor B cells of affinity greater than 3*µM* were considered similar to earlier works^10^. Accordingly, 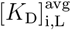, which was equivalent to 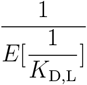, was calculated as 0.195*µM*. For all cases, 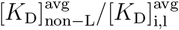 was rounded to the nearest order of magnitude.

The error factor *ϵ* in Equation 1 was estimated using a singular reference data point (obtained at day 8 post immunization) from the experiments of Huang et al.^17^, i.e., where bnAb precursor affinity was intermediate and frequency was 1 in 10^3^. To do this, we exploited the observation that for GC simulations performed with different values of 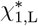 close to the reference data point, the percentage of bnAb-type GC B cells evolved almost linearly (Fig.S1A). Thus, by linearly extrapolating the reference data point to the time of GC onset, 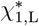 was calculated as 0.02318, which in turn allowed estimation of *ϵ* as 0.0255.

#### GC Simulation Model

The GC mathematical model used here is adapted from earlier versions with modifications^57,64,78,81^ A comprehensive description is presented below. Used acronyms are: DZ for dark zone, LZ for light zone, Tfh for T follicular helper cell, FDC for follicular dendritic cell.

#### Space representation

All reactions take place on a three-dimensional discretized space with a rectangular lattice with lattice constant of Δ*x* = 5*µm*. Every lattice node can be occupied by a single cell only.

#### Shape space for antibodies

Antibodies are represented on a four (*d* = 4) dimensional shape space^109^. The shape space is restricted to a size of 10 positions per dimension, thus only considering antibodies with a minimum affinity to the antigen. The optimal clone *φ*^∗^ is positioned in the centre of the shape space. A position on the shape space *φ* is attributed to each B cell. The 1-Norm with respect to the optimal clone 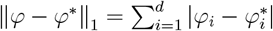, i.e., the minimum number of mutations required to reach it (mutation distance), is used as a measure for the antigen binding probability. The binding probability is calculated from the Gaussian distribution with width Γ = 2.8^63^:

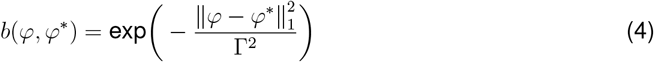

#### B cell phenotypes

Three B cell phenotypes are distinguished: DZ B cells, LZ B cells, and plasma cells. The different phenotypes characterize the cell properties and are not meant as localization within the GC zones. DZ B cells divide, mutate, and migrate. LZ B cells also migrate and undergo the different stages of the selection process. Plasma cells migrate and produce antibodies.

#### GC initialization

The model starts from 200 Tfh, 200 FDCs, 300 stromal cells, and no B cell. Tfh are randomly distributed on the lattice and occupy a single node each. Stromal cells are restricted to the DZ (see section Chemokine distribution for their function). FDCs are restricted to the upper half of the reaction sphere, occupy one node by their soma and have 6 dendrites of 40*µm* length each. The presence of dendrites is represented as a lattice-node property and, thus visible to B cells. The dendrites are treated as transparent for B cell or Tfh migration such that they do not inhibit cell motility.

Next, GCs were initialized by the entry of B founder cells (calculated in Pre-GC Model). These cells enter the GC space at a rate of 2 cells per hour (see Calculation of B cell influx rate) till 96 hours, resulting in about 180 - 200 founder cells^110^, a value which is consistent with experiments^77^. The amount of available antigen in lymph nodes and the number of available Tfh cells in the pre-GC phase can influence founder cell activation^40^. However, as only single bolus dose immunization in the context of HIV immunogens was considered, these variables and consequently the total number of founder cells were assumed to be invariable across simulations.

Depending on the composition calculated in Equation 3, founder cells were tagged as bnAb- or non-bnAb-type and mapped to appropriate mutation distances (ranging from 8 to 4) based on their average *K*_D_ values (ranging from 50 to 0.1*µM*). To do this, five *K*_D_ intervals were defined as [50, 10[, [10, 5[, [5, 1[, [1, 0.5[and [0.5, 0.1[. As the first *K*_D_ interval had the lowest affinity, it was assumed to have the largest mutation distance of 8. For each successive *K*_D_ interval, the associated mutation distance decreased by one. All founder cells were assumed to follow identical selection rules, irrespective of type or lineage within the GC. Masking by antibodies derived from GC-exiting plasma cells was always present (unless specified otherwise).

#### Calculation of B cell influx rate

As B cell selection is not active during the first 3 days of the reaction (i.e., days 3-5 post immunisation), the first 3 days can be approximated as a B cell expansion phase. Clonality can be safely ignored, and it suffices to consider a single dividing cell type Bi, where i denotes the generation number of the B cells. The dynamics of expansion are then described by

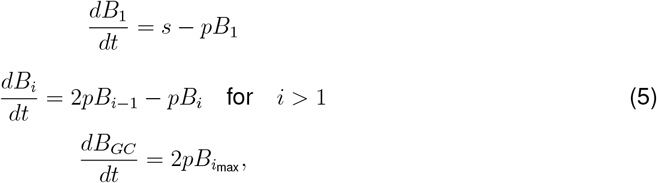

where *s* is the influx rate, *p* the division rate, *i*_max_ the number of divisions per cell in the expansion phase, and *B*_*GC*_ the resulting number of GC B cells that participate in the GC reaction. The number of initial divisions is estimated by the maximum number of divisions observed upon anti-DEC205-OVA treatment, i.e., *i*_max_ = 6^45,64^. As the division time ln(2)*/p* is shorter than the expansion phase *T*_expand_ = 3 days, one may solve Eq.5 in steady state, yielding:

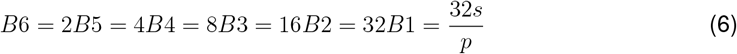

Thus, the relevant ODE becomes

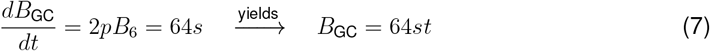

i.e., a linear growth in time proportional to the constant influx during expansion. Note that in the steady state approximation the influx rate becomes independent of the division rate *p*, which is an implication of the assumption of a fixed number of divisions per founder cell *i*_max_. With the side condition of getting 9000 cells at day 3, *B*_GC_(*T*_expand_ = 72hr) = 9000, the influx rate is estimated to be

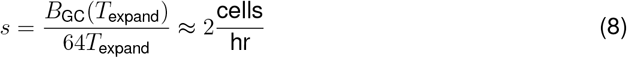

This corresponds to 144 B cells entering the GC in the first 3 days of expansion and building up the founder cell population of the GC reaction. Motivated by this estimation, in the model we assumed that B cells enter the GC reaction with a probability corresponding to a rate of 2 cells per hour for 96 hours. New B cells are randomly positioned on the lattice (exclusively on free nodes).

#### Antigen-presentation by FDCs

Each FDC is loaded with 3500 antigen portions distributed onto the lattice nodes occupied by FDC-soma or FDC-dendrite. One antigen portion corresponds to the number of antigen molecules taken up by a B cell upon successful contact with an FDC.

For multiple epitopes, a single antigen with multiple epitopes is considered. Here, antigenic degradation was assumed to be minimal such that each antigen deposit on the FDC had all its epitopes intact. Therefore, all epitopes were presented in equal stoichiometry on the FDC. Additionally, B cells could take up the entire antigen upon successful interaction, following which, all epitopes displayed on the relevant FDC site were decreased by one. All these are important changes from^57^

For all scenarios with two epitopes simulated here, the epitopes were assumed to be structurally dissimilar to minimize B cell crossreactivity for the two epitopes. In addition, it was assumed that the two epitopes were spatially separated such that the AEM directed towards one epitope did not alter B cell binding to the second epitope. To implement these two assumptions, the positions of the two epitopes were chosen sufficiently far apart, i.e., at positions 3333 and 6666, respectively, of the shape space grid (see Shape space for antibodies).

#### Antigen-antibody interaction on FDCs

Antibodies in this work are mainly sourced from plasma cells of the GC reaction (see section Antibody sources below) and are represented in the 4-dimensional shape space with 10 positions in each direction. The quantity of interest is the amount of free antigen at each FDC site when antibodies are present and changing over time. As it is not feasible to calculate the amount of free antigen at each site for all 10,000 possible antibody types, 11 bins, *B*_ij_, with *i* element of [0, …, 10] based on antibody-binding-probability were introduced, where the binding probability was defined relative to an antigen-epitope *j*. Thus every epitope gives rise to its own antibody distribution on the bins. We assumed a constant on-rate *k*_on_ = 10^6^*/*(Mol sec)^111^ and a variable off-rate

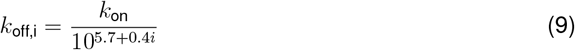

mimicking a dissociation constant that varies over 4 orders of magnitude. At each FDC site *x*, the chemical kinetics equation for the immune complexes *C*_ij_(*x*) formed between antibodies in bin *i* and epitope *j*

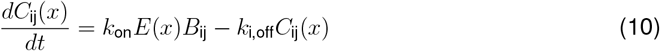

was solved for every epitope *j* in order to determine the amount of free antigen *E*(*x*) at this site. Only this amount of antigen is available for B cells to bind antigen with probability according to Eq.4. In addition to the above, competitive binding between B cells and antibodies for the calculated free epitopes is assumed^54^. This is implemented by scaling down the binding probability of B cells in Eq.4 as

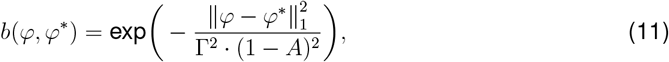

where *A* is the average binding probability of all antibodies present.

#### DZ B cell division

The average cell cycle duration of 7 h of DZ B cells is varied for each B cell according to a Gaussian distribution. This is needed to get desynchronization of B cell division. The cell cycle is decomposed into four phases (G1, S, G2, M) in order to localize mitotic events if this is needed. Each founder cell divides a number of times before differentiating to the LZ phenotype for the first time. Six divisions was the number of divisions found in response to the extreme stimulus with anti-DEC205-OVA^45,64^. Each selected B cell divides a number of times determined by the interaction with Tfh (see below, LZ B cell selection). The parameters of the interaction with Tfh are tuned such that the mean number of divisions is in the range of two^86^. This value is required in order to maintain a DZ to LZ ratio in the range of two^45,64^. A division requires free space on one of the Moore neighbors of the dividing cell. Otherwise the division is postponed until a free Moore neighbor is available. At every division the encoded antibody can mutate with a probability of 0.5^112,113^. This corresponds to a shift in the shape space to a von Neumann neighbor in a random direction. Upon selection by Tfh the mutation probability is individually reduced from *m*_max_ = 0.5 down to *m*_min_ = 0 in an affinity-dependent way following

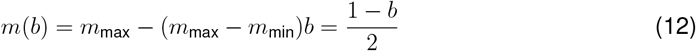

with b from Eq.4^114^. Thus, after recycling DZ B cells can acquire reduced mutation probabilities. This mechanism is motivated by the observation that B cell receptor internalization enhances the activation of the kinase Akt^115^ which, in turn, suppresses activation induced cytosine deaminase (AID)^116^. AID is required for somatic hypermutation, such that this provides an affinity dependent downregulation of the mutation frequency^117^. Although, there is no formal proof of this mechanism, a similarly downregulation of AID was observed for B cells undergoing a high number of proliferations (clonal bursts) in the DZ^118^. B cell division of B cells that previously acquired antigen and have been selected by Tfh distribute the retained antigen asymmetrically to the daughters^119^. The model assumes asymmetric division in 72% of the cases, which is supported by experimental observations (see^119^ and Figure S1 in^64^). If division is asymmetric, one daughter gets all the retained antigen while the other gets none, which approximates the value of 88% found in^119^. Mutation is suppressed in B cells with retained antigen. After the required number of divisions, the B cell differentiates with a rate of in 1/6 min to the LZ phenotype. All B cells that kept the antigen up to this time, differentiate to plasma cells, upregulate CXCR4, and leave the GC in direction of the T zone.

#### LZ B cell selection

LZ B cells can be in the states unselected, FDC-contact, FDC-selected, Tfh-contact, selected, apoptotic.

#### Unselected

LZ B cells migrate and search for contact with FDCs loaded with antigen in order to collect antigen for 0.7 h. If an FDC soma or dendrite is present at the position of the B cell, the B cell attempts to establish contact to the epitope with highest affinity to the B cell receptor. Binding is affinity dependent and happens with the probability *b* in Eq.4. If the available number of antigen portions at the specific FDC site drops below 20 the binding probability b is linearly reduced with the number of available portions. If successful, the B cell switches to the state *FDC-contact*; otherwise the B cell continues to migrate. Further binding-attempts are prohibited for 1.2 min. At the end of the antigen collection period, B cells switch to the state *FDC-selected*. If a LZ B cell fails to collect any antigen at this time it switches to the state *apoptotic*.

#### FDC-contact

LZ B cells remain immobile (bound) for 3 min^84^ and then return to the state *unselected*. The counter for the number of successful antigen uptake events is increased by one and the FDC reduces its locally available antigen portions by one.

#### FDC-selected

B cells search for contact with Tfh. If they meet a Tfh they switch to the state *Tfh-contact*.

#### Tfh-contact

LZ B cells remain immobile for 6 min. In this time the bound Tfh, which may also be bound to other B cells, polarizes to the bound B cell with highest number of successful antigen uptakes. Only this B cell receives Tfh signals and accumulate those. After the binding time, the B cell detaches and returns to the state *FDC-selected*. It continues to search and bind Tfh cells until the Tfh search time of 3 h is over. Then, it switches to the state *apoptotic* if the accumulated Tfh-signaling time remained below 30 min. Otherwise it switches to the state *selected*.

#### Selected

LZ B cells keep the LZ phenotype for six hours and desensitize for CXCL13, thus perform a random walk. During that time they re-enter cell cycle and progress through the cell cycle phases. Then they recycle back to the DZ phenotype with a rate of 1/6 min and memorize the amount of collected antigen as well as the cell cycle phase they have achieved by this time.

The number of divisions *P* (*A*) the recycled BCs will do is derived from the amount of collected antigen *A*, which reflects the amount of pMHC presented to Tfh and the affinity of the BCR for the antigen, according to

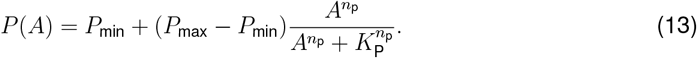

The minimum number of division is set to one *P*_min_ = 1 in order to avoid recycling events without further division. It is limited by 6 divisions in the best case, which is motivated by anti-DEC205-OVA experiments in which DEC205+*/*+ BCs received abundant antigen which increased pMHC presentation to a maximum^45^. The population dynamics *in vivo* and *in silico* only matched when the number of divisions was increased to six in the simulation^64^ suggesting that the strongest possible pMHC presentation to Tfh cells induces six divisions *P*_max_ = 6. The Hill-coefficient *n*_p_ = 1.8.

The half value *K*_P_ remained to be determined, which denotes the amount of antigen collected by B cells at which the number of divisions becomes half maximal. The number of collected antigen portions varies between zero and a maximum determined by the duration of the antigen collection phase, the duration of each B cell interaction with FDCs, and the migration time between two antigen presenting sites. The numbers of successful B cell-FDC encounters as observed in the simulations served as estimate of Amax. For an intermediate antigen uptake of *A*_0_ = 3, the resulting number of divisions has to be *P*_0_ = 2 in order to be in agreement with the mean number of divisions in the range of two^86^, which leads to the condition:

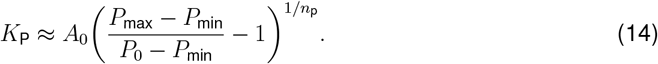

#### Apoptotic

LZ BCs remain on the lattice for 6 hours before they are deleted. They continue to be sensitive to CXCL13 during this time.

### Antibody sources

#### Plasma cells

Plasma cells from the GC reaction are collected and memorized together with the affinity of the encoded antibody to the epitopes. Their life time is assumed longer than the duration of the GC reaction. Plasma cells are attributed to the different affinity bins for each epitope and further differentiate to an antibody forming plasma cell according to a linear rate equation with a rate of ln(2) per day, i.e., with a half life of one day. All plasma cells produce antibodies *B*_ij_, attributed to bin *i* for each epitope *j*

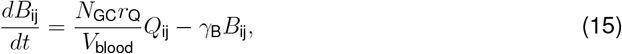

where *r*_Q_ = 3 · 10^18^mol/hour is the production rate per cell^120^ and *γ*_B_ = ln(2)/(14days) is the degradation rate of antibodies. The produced antibodies are assumed to distribute over the whole organism. The sum of all *N*_GC_ GC reactions in the organism is diluted over the mouse blood volume *V*_blood_ and adds up to the total antibody concentration found in the simulated GC. The factor *N*_GC_*r*_Q_*/V*_blood_ = 25*/*ml was assumed. This setting implies that antibodies are homogeneously distributed over the space of the simulated GC. An alternative source of antibodies comes from their injection.

### Chemokine distribution

Two chemokines CXCL12 and CXCL13 are considered. CXCL13 is produced by FDCs in the LZ with 10nMol per hour and FDC while CXCL12 is produced by stromal cells in the DZ with 400nMol per hour and stromal cell. As both cell types are assumed to be immobile, chemokine distributions were pre-calculated once and the resulting steady state distributions were used in all simulations.

### Chemotaxis

DZ and LZ BCs regulate their sensitivity to CXCL13 and CXCL12, respectively. This is true in all BC states unless stated otherwise. All BCs move with a target speed of 7.5*µm/*min. This leads to a slightly lower observable average speed of ≈ 6*µm/*min. BCs have a polarity vector that determines their preferential direction of migration. The polarity vector 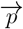 is reset every 1.5 minutes into a new direction using the chemokine distribution *c* as

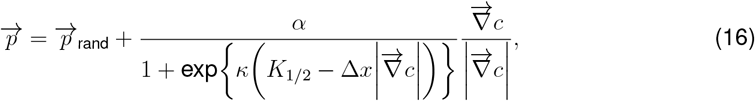

where 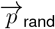 is a random polarity vector and the turning angle is sampled from the measured turning angle distribution (^82^, Figure S1B). *α* = 3 determines the relative weight of the chemotaxis and random walk, *K*_1*/*2_ = 2 · 10^11^ Mol determines the gradient of half maximum chemotaxis weight, and *κ* = 10^10^*/*Mol determines the steepness of the weight increase.

BCs de- and re-sensitize for their respective chemokine depending on the local chemokine concentration: The desensitization threshold is set to 4.5 nMol and 0.08 nMol for CXCL12 and CXCL13, respectively, which avoids cell clustering in the centre of the zones. The resensitization threshold is set at 2/3 and 3/4 of the desensitization threshold for CXCL12 and CXCL13, respectively. BCs can only migrate if the target node is free. If occupied and the neighbor cell is to migrate in the opposite direction (negative scalar product of the polarity vectors) both cells are exchanged with a probability of 0.5. This exchange algorithm avoids lattice artifacts leading to cell clusters. Tfh cells do random walk with a preferential directionality to the LZ: The polarity vector 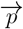 of Tfh cells is determined from a mixture of random walk 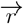 and the direction of the LZ 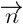 by

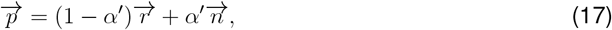

where *α*^′^ = 0.1 is the weight of chemotaxis. This weight leads to a dominance of random walk with a tendency to accumulate in the LZ as found in experiment. Tfh cells migrate with an average speed of 10*µm/*min and repolarize every 1.7 minutes^121^. Plasma cell motility is derived from plasma cell motility data to 3*µm/*min with a persistence time of 0.75 minutes^82^.

### Simulation formulation for arbitrary number of bnAb lineages

To evaluate a generalized setting involving an arbitrary number of bnAb lineages the following setup was used.

For simultaneous priming (scenario (i)), bnAb precursor B cells were aggregated based on their epitope specificity. Overall responses, measured as the AUC of number of bnAb-type GC B cells, *AUC*_sc.1_, could now be calculated using total epitope-specific bnAb precursor frequencies in Equations 1 and 3 and subsequently simulating GCs. This was reasonable as only high affinity bnAb precursor B cells 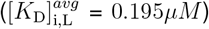 were considered since associated responses are most relevant for GC reactions^16–18^.

For scenario (ii), simulations were performed by splitting the overall targeted bnAb precursor pool into smaller subsets. These subsets were ubiquitously defined in the lower frequency limit (*B*_0,L_ = 1 in 10^7^), had specificity entirely directed towards one of the two epitopes and in consistency with scenario (i) were of high affinity. Accordingly, response of a bnAb precursor pool with overall frequency as 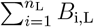, can be approximated by 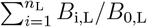 iterations of the above defined precursor subset. In other words, the total lineage AUC for scenario (ii), *AUC*_sc.2_, can be written as

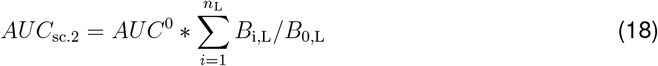

where *AUC*^0^ is the AUC of a singular bnAb subset.

To compare the two scenarios, additivity index, Ω, was defined as

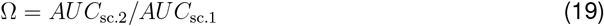

### Quantification and statistical analysis

For every parameter set we simulated 400 GCs to obtain sufficient statistics. As a single lymph node can have multiple GCs, to simulate the behaviour of a lymph node we distributed these runs into sets of *m* = 4 with *n* = 100 GCs each. For each set, the means of various quantities were calculated. These set means were used to calculate overall means and standard deviations as described below.

#### Affinity of GC B cells

The average affinity of GC B cells was calculated using binding probability *b* at time *t* as

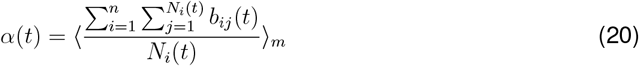

where *b*_*ij*_(*t*) was the affinity of the *j*^th^ B cell among *N*_*i*_(*t*) B cells of the *i*^*th*^ GC of a set.

#### Number of GC B cells

The average number of GC B cells at time *t* was calculated as

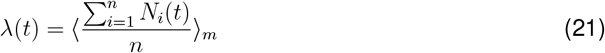

#### Percentage of lineage-specific GC B cells

The average percentage of GC B cells belonging to a specific bnAb lineage at time *t* was calculated as

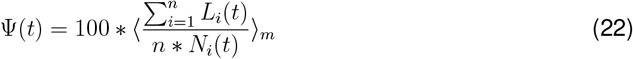

where *L*_*i*_(*t*) was the total number of lineage-specific GC B cells in the *i*^th^ GC of a set.

