## Supplementary figures for "Multiple broadly neutralizing antibody lineages can co-exist and mature in the same germinal centres"

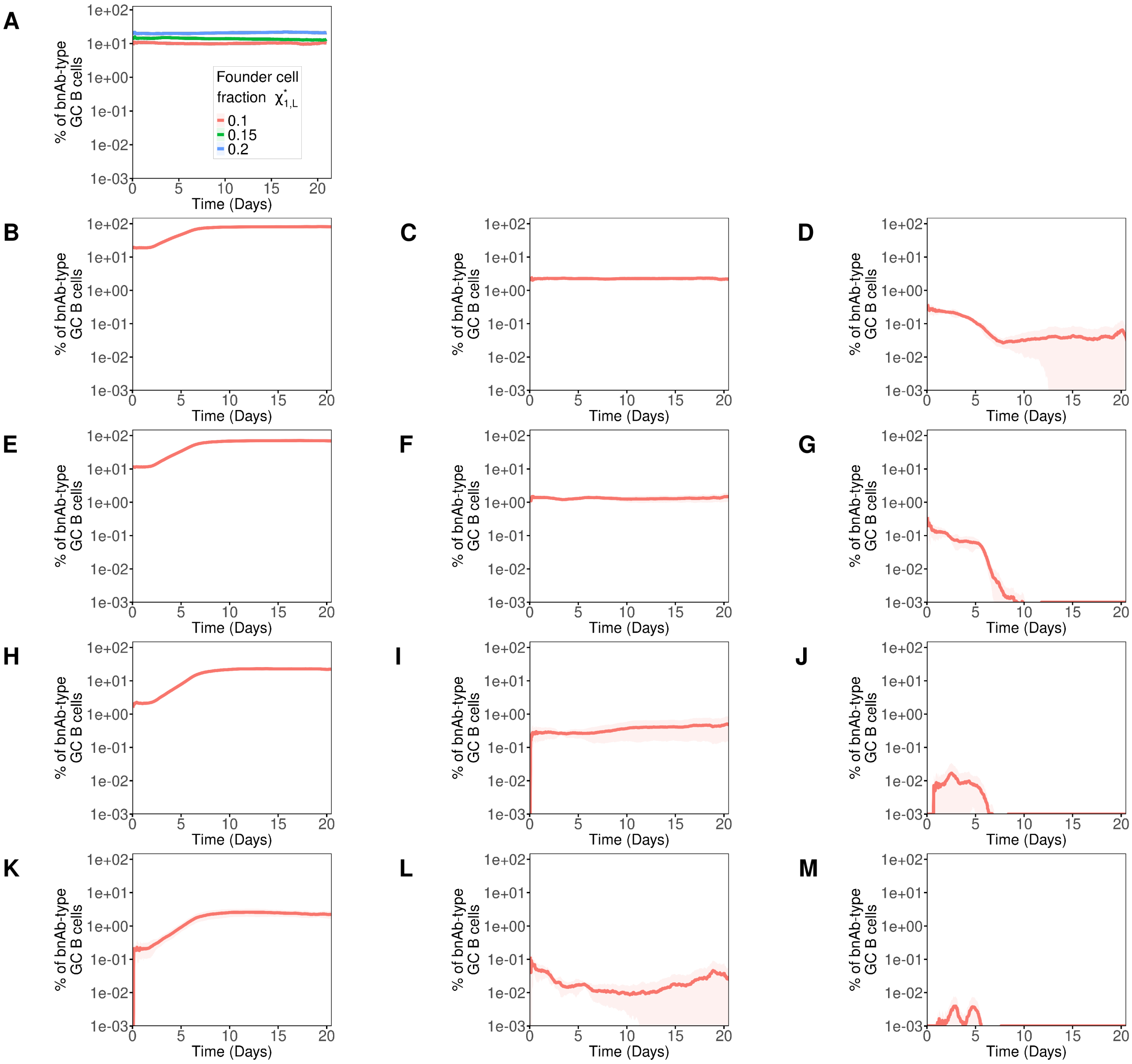


Figure S1: **Temporal bnAb lineage responses in relation to Fig.2.** (A) Percentage of bnAb-type GC B cells for intermediate bnAb precursor affinity with $\chi^*_{1,\mathrm{L}}$ fixed as 0.1, 0.15 and 0.2. (B-M) percentage of bnAb-type GC B cells for different precursor frequencies and affinities. Precursor frequency is varied row-wise as (B-D) $1$ in $10^3$, (E-G) $1$ in $10^4$, (H-J) $1$ in $10^5$ and (K-M) $1$ in $10^6$. Precursor affinity is varied column-wise as (B, E, H and K) high, (C, F, I and L) intermediate and (D, G, J and M) low. Means (lines) were calculated by averaging the pooled means obtained from 4 sets of 100 simulations. Standard deviation (shaded area) is calculated as the standard error of means.

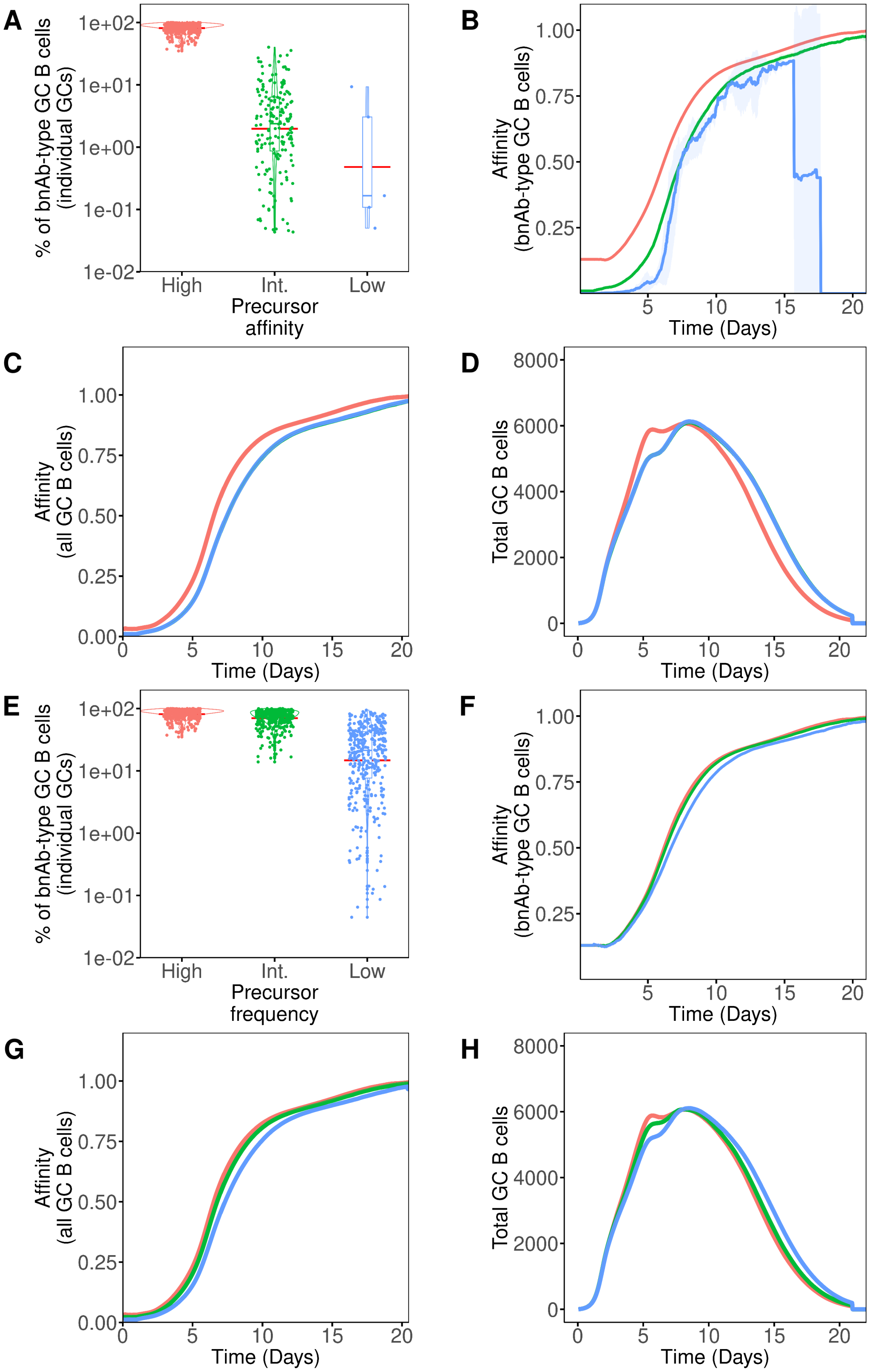


Figure S2: **Temporal bnAb lineage responses in relation to Fig.2.** Temporal trends of GCs simulated with low (blue), intermediate (green) and high (red) (A-D) bnAb precursor affinities and (E-H) frequencies are as follows: (A, E) percentage of bnAb-type GC B cells in individual GCs at day 17 of GC reaction, (B, F) average affinity of bnAb-type GC B cells (only GCs with 2 or more bnAb-type cells were considered) (C, G) average affinity of all GC B cells, (D, H) averaged total GC B cell count. Means (lines) were calculated by averaging the pooled means obtained from 4 sets of 100 simulations. Standard deviation (shaded area) was calculated as the standard error of means. The violin plots show the median (horizontal line inside the box plot), 25 and 75 percentiles and the mean (horizontal red line).

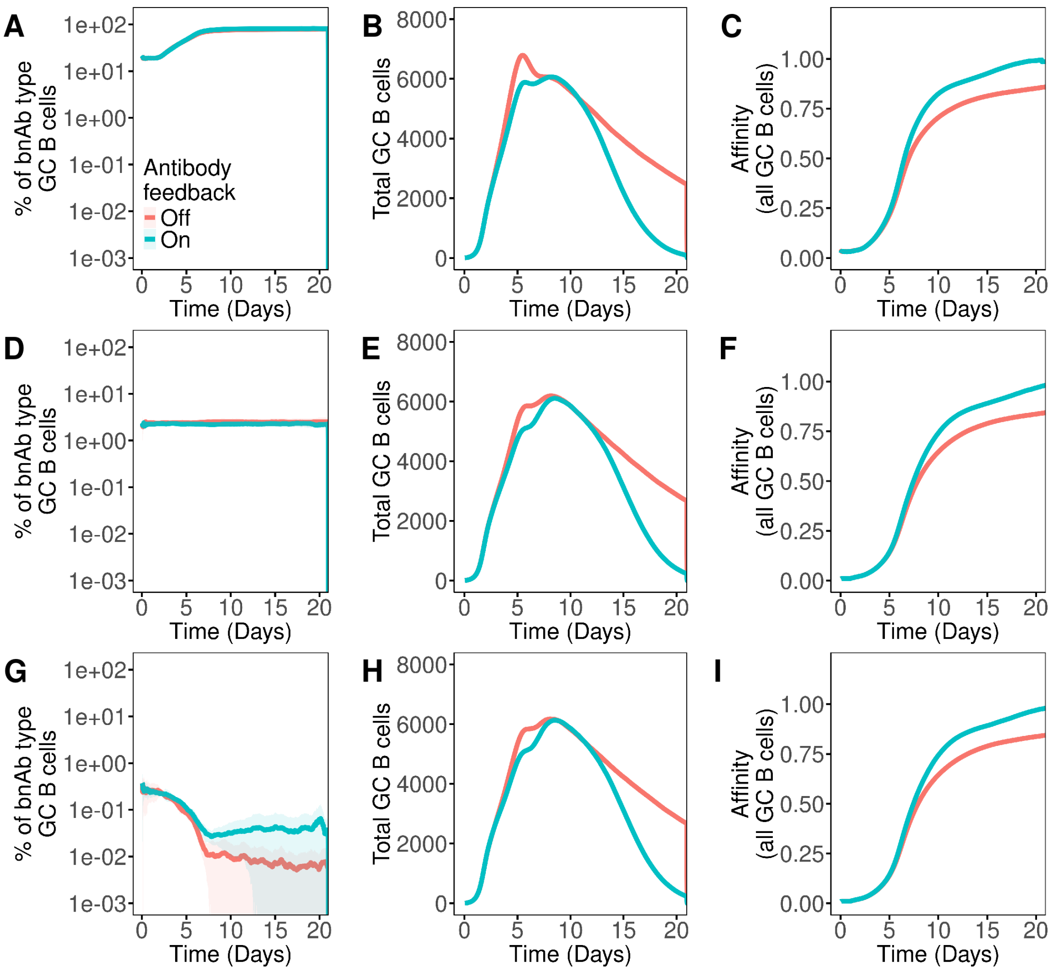


Figure S3: **Impact of AEM on bnAb lineage evolution in single-epitope scenario.** BnAb lineage was simulated in the presence (blue) or absence of AEM (red) and it's precursor frequency was fixed as $1$ in $10^3$ while precursor affinity ($K_{\mathrm{D}}$) was varied as (A-C) high ($0.125\mu M$), (D-F) intermediate ($1.3\mu M$) or (G-I) low ($18.5\mu M$). Temporal trends of GCs as: (A, D and G) percentage of bnAb-type GC B cells, (B, E and H) averaged total GC B cell count and (C, F and I) average affinity of all GC B cells. Means (lines) were calculated by averaging the pooled means obtained from 2 sets of 50 simulations. Standard deviation (shaded area) was calculated as the standard error of means.

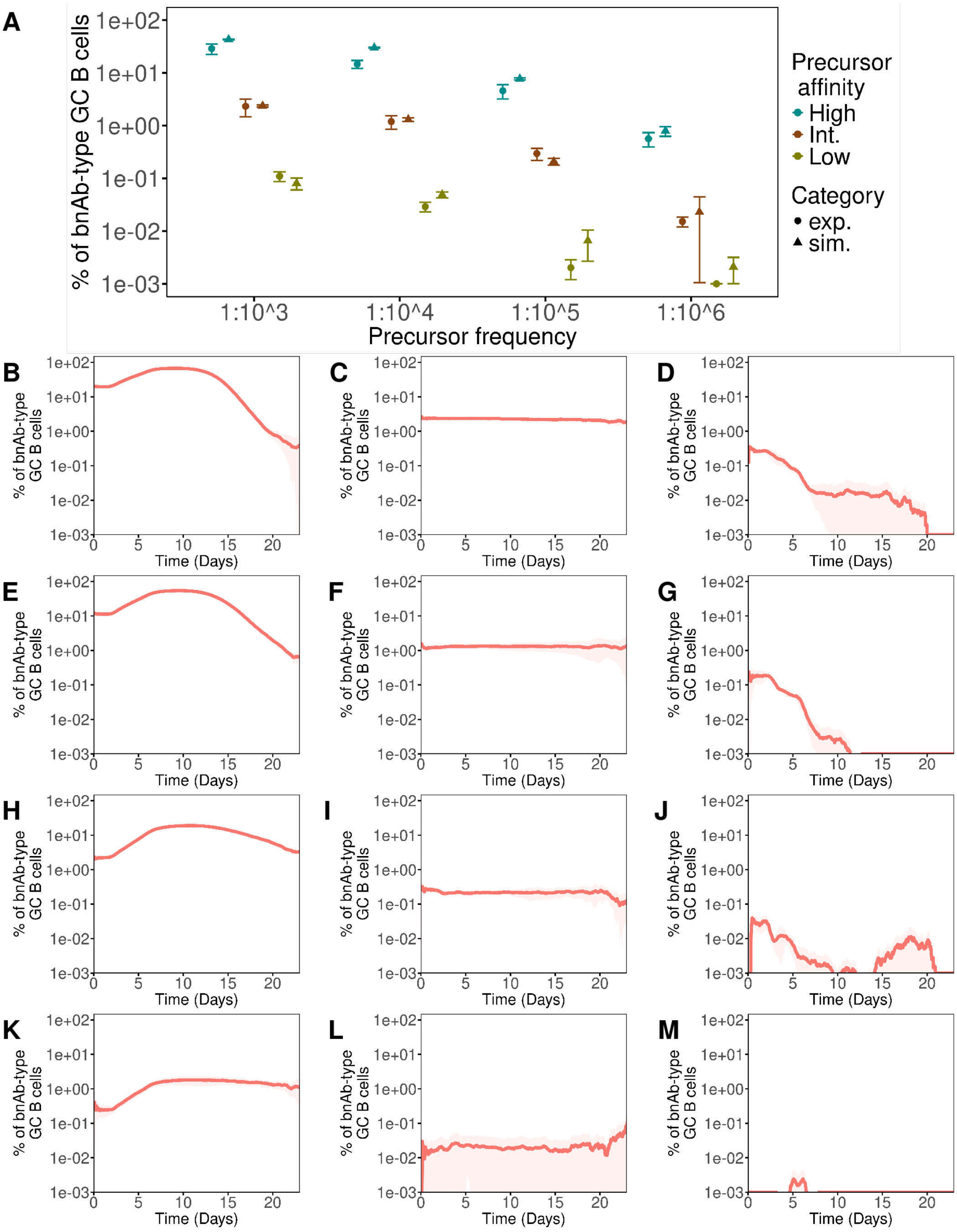


Figure S4: **Multi-epitope simulations recapitulate the precursor effect as seen in experiments.** (A) Comparison of bnAb-type GC B cells in experiments[1] and simulations. BnAb precursor affinity ($K_{\mathrm{D}}$) in wildtype C57BL/6 mouse was varied as high ($0.125\mu M$), intermediate ($1.3\mu M$) or low ($18.5\mu M$) while the frequency was 10-fold serially diluted from $1$ in $10^3$ to $1$ in $10^6$. Accounting for the time taken for GC onset[2], simulation data (triangles) at day 5 post GC onset was compared to experimental data (circles) at day 8 post immunization. (B-M) Percentage of bnAb-type GC B cells for different precursor frequencies and affinities. Precursor frequency is varied row-wise as (B-D) $1$ in $10^3$, (E-G) $1$ in $10^4$, (H-J) $1$ in $10^5$ and (K-M) $1$ in $10^6$. Precursor affinity is varied column-wise as (B, E, H and K) high, (C, F, I and L) intermediate and (D, G, J and M) low. Means (lines and points) were calculated by averaging the pooled means obtained from 4 sets of 100 simulations. Standard deviation (error bars and shaded areas) is calculated as the standard error of means.

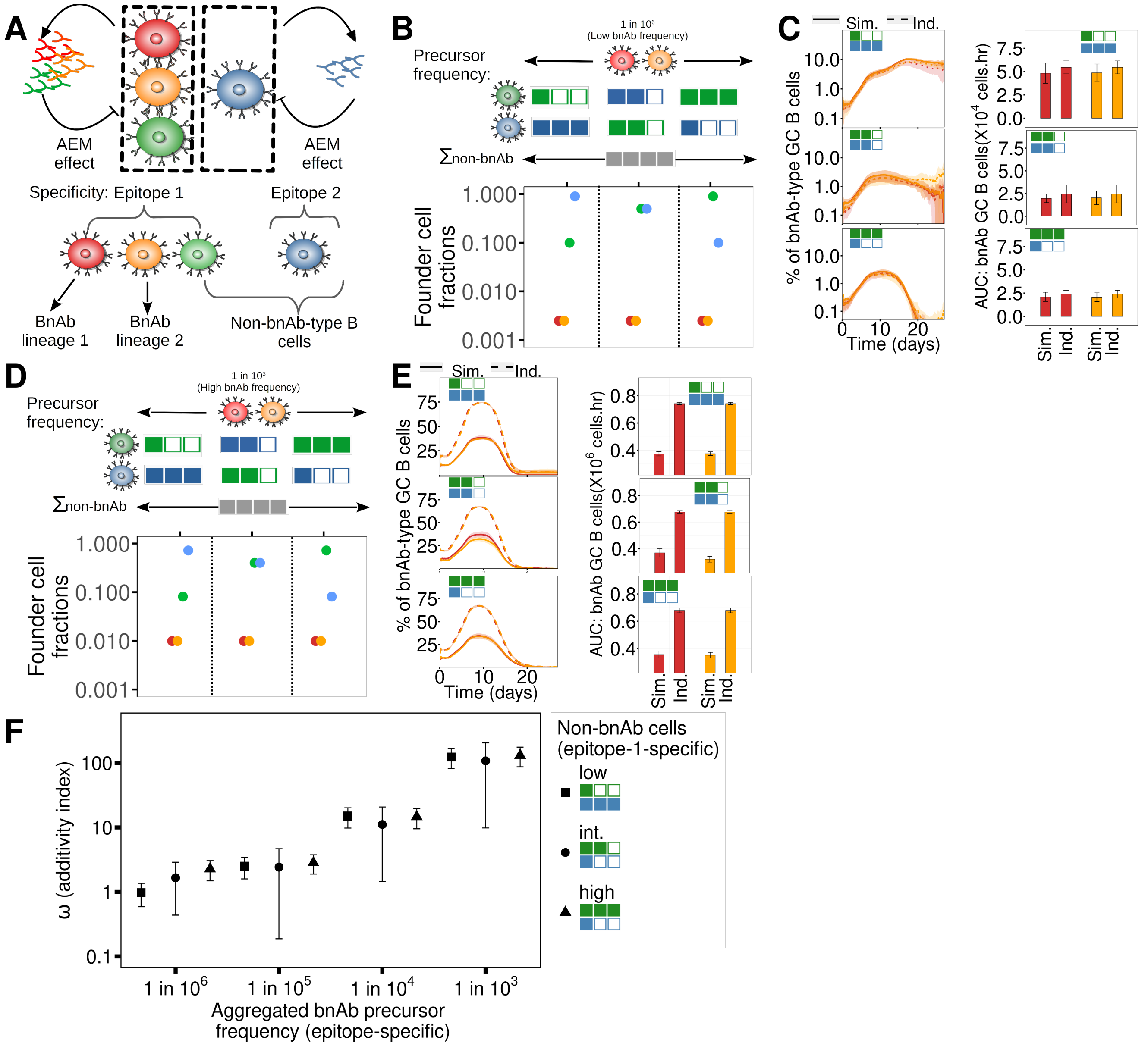

Figure S5: **Simultaneous vs. individual priming of multiple bnAb lineages with shared epitope specificities.** (A) Simulation schematic for multiple lineages with AEM. For simultaneous priming, bnAb lineages are simulated concurrently in a single GC ensemble, while for individual. (B-F) $90\%$, $50\%$ or $10\%$ non-bnAb precursors were specific for epitope $1$. For both bnAb lineage $1$ (red) and $2$ (orange), harmonic mean precursor affinity was high ($0.195\mu M$) while frequency was low (B, C, $1$ in $10^6$) or high (D, E, $1$ in $10^3$). GC B cell characteristics: (B, D) founder cell fractions, (C, E left) percentage of bnAb-type cells for simultaneous (solid lines) or individual (dashed lines) lineage priming and (C, E right) AUC of bnAb-type cells. (F) $\Omega$ (additivity index) calculated for an arbitrary number of bnAb lineages. Colored boxes represent classes of the frequencies of non-bnAb precursor B cells and do not reflect actual values or scale. Means (bars and lines) were calculated by averaging the pooled means obtained from 4 sets of 100 simulations. Standard deviation (shaded area and error bars) was calculated as the standard error of means.
